# *Pseudomonas aeruginosa* surface motility and invasion into competing communities enhances interspecies antagonism

**DOI:** 10.1101/2024.04.03.588010

**Authors:** Andrea Sánchez-Peña, James B. Winans, Carey D. Nadell, Dominique H. Limoli

## Abstract

Chronic polymicrobial infections involving *Pseudomonas aeruginosa* and *Staphylococcus aureus* are prevalent, difficult to eradicate, and associated with poor health outcomes. Therefore, understanding interactions between these pathogens is important to inform improved treatment development. We previously demonstrated that *P. aeruginosa* is attracted to *S. aureus* using type IV pili-mediated chemotaxis, but the impact of attraction on *S. aureus* growth and physiology remained unknown. Using live single-cell confocal imaging to visualize microcolony structure, spatial organization, and survival of *S. aureus* during coculture, we found that interspecies chemotaxis provides *P. aeruginosa* a competitive advantage by promoting invasion into and disruption of *S. aureus* microcolonies. This behavior renders *S. aureus* susceptible to *P. aeruginosa* antimicrobials. Conversely, in the absence of type IV pilus motility, *P. aeruginosa* cells exhibit reduced invasion of *S. aureus* colonies. Instead, *P. aeruginosa* builds a cellular barrier adjacent to *S. aureus* and secretes diffusible, bacteriostatic antimicrobials like 2-heptyl-4-hydroxyquinoline-*N*-oxide (HQNO) into the *S. aureus* colonies. *P. aeruginosa* reduced invasion leads to the formation of denser and thicker *S. aureus* colonies with significantly increased HQNO-mediated lactic acid fermentation, a physiological change that could complicate the effective treatment of infections. Finally, we show that *P. aeruginosa* motility modifications of spatial structure enhance competition against *S. aureus*. Overall, these studies build on our understanding of how *P. aeruginosa* type IV pili-mediated interspecies chemotaxis mediates polymicrobial interactions, highlighting the importance of spatial positioning in mixed-species communities.

## IMPORTANCE

The polymicrobial nature of many chronic infections makes their eradication challenging. Particularly, coisolation of *Pseudomonas aeruginosa* and *Staphylococcus aureus* from airways of people with cystic fibrosis and chronic wound infections is common and associated with severe clinical outcomes. The complex interplay between these pathogens is not fully understood, highlighting the need for continued research to improve management of chronic infections. Our study unveils that *P. aeruginosa* is attracted to *S. aureus*, invades into neighboring colonies, and secretes anti-staphylococcal factors into the interior of the colony. Upon inhibition of *P. aeruginosa* motility, and thus invasion, *S. aureus* colony architecture changes dramatically, whereby *S. aureus* is not only protected from *P. aeruginosa* antagonism, but also responds through physiological alterations that may further hamper treatment. These studies reinforce accumulating evidence that spatial structuring can dictate community resilience and reveal that interspecies chemotaxis and bacterial motility are critical drivers of interspecies competition.

## INTRODUCTION

Many microorganisms exist in complex polymicrobial environments, such as the soil and the human body, where they interact with each other and respond to changes in their surroundings [1–3]. These interactions can lead to the emergence of community-level properties, which may not be observed in monoculture [4–6]. The resulting collective behavior can have significant implications for the physiology and evolution of microbial populations and their interactions with the host. For example, bacterial pathogens can enhance both virulence and antibiotic tolerance in mixed communities [2, 4, 7–12], which may contribute to the ineffectiveness of current therapeutic strategies against chronic infections.

*Pseudomonas aeruginosa* and *Staphylococcus aureus* are the most prevalent and abundant pathogens in individuals with cystic fibrosis (CF) [5, 13] and can coexist and persist in significant quantities in the lungs for decades [14]. Critically, their coinfection is linked with more severe lung disease, increased rates of hospitalization, and reduced lung function in patients [15–19]. Additionally, coinfection in chronic burn wounds can delay healing time [20]. Thus, there is a need to further understand how interactions between these two organisms can worsen the outcomes of polymicrobial infections.

Several *in vitro* studies support clinical observations that *P. aeruginosa* and *S. aureus* increase each other’s virulence during coculture [8, 21]. *P. aeruginosa* secretes an arsenal of anti-staphylococcal factors including the siderophores pyoverdine and pyochelin, phenazines, rhamnolipids, staphylolytic proteases like LasA, and quinolones [22–28]. Many of these secreted products alter *S. aureus* physiology in a manner that increases its tolerance to antibiotics [9, 11, 12, 29–31]. One example is the small molecule, 2-heptyl-4-hydroxyquinoline *N*-oxide (HQNO), which inhibits *S. aureus* cellular respiration, and in turn, can shift *S. aureus* metabolism from respiration to fermentation [7, 26, 32, 33]. HQNO has been detected in sputum from people with CF [26], and its mechanism of action allows for an increase in *S. aureus* tolerance to several antibiotics used clinically to treat *S. aureus* infections in CF [9, 11, 12]. However, the effect of these antimicrobials on *S. aureus* has been mainly studied by adding cell-free supernatants of *P. aeruginosa* to *S. aureus* in a well-mixed environment [9, 11, 12, 31, 34]. Interestingly, recent studies found that HQNO modifies the spatial organization of *P. aeruginosa* and *S. aureus* in a synthetic CF sputum medium (SCFM2) [35] and *in vivo* in chronic murine wounds [36], which highlights the importance of visualizing these microorganisms together in a structured environment. Therefore, to further elucidate how interspecies interactions negatively impact clinical outcomes, there is a need to study these organisms in an experimental model system that better reflects how microbes naturally encounter each other, namely, under spatial constraint.

Previously, we demonstrated that *P. aeruginosa* responds to *S. aureus* from a distance by increasing type IV pili motility, retractile appendages that allow *P. aeruginosa* to move across surfaces in a mode of locomotion referred to as twitching motility [37]. By beginning experiments where *P. aeruginosa* and *S. aureus* are spatially separate as single cells, allowing each species to grow and reciprocally respond to each other at a distance, we observed that *P. aeruginosa* directionally moves towards *S. aureus* and engages with the surface of *S. aureus* microcolonies [38]. *P. aeruginosa* attraction towards *S. aureus* occurs through detection of *S. aureus* secreted peptides called phenol-soluble modulins (PSMs) by the Pil-Chp chemoreceptor, PilJ [39, 40]. However, it remains unknown how *P. aeruginosa* chemotaxis towards *S. aureus* influences *S. aureus* physiology and survival.

Here we sought to understand the consequences of *P. aeruginosa* type IV pili-mediated interspecies attraction on *S. aureus*. We visualized the course of *P. aeruginosa* and *S. aureus* interactions at the single-cell level over time using resonant scanning confocal microscopy and discovered that *P. aeruginosa* utilizes a combination of type IV pili-mediated motility and secreted antimicrobials to effectively outcompete *S. aureus* under these conditions. Particularly, we found that wild-type (WT) *P. aeruginosa* is attracted to, invades, and disrupts *S. aureus* colonies. Moreover, *P. aeruginosa*-secreted antimicrobials HQNO, pyoverdine, pyochelin, and LasA were necessary for negatively influencing *S. aureus* growth, but not for invasion and disruption of *S. aureus* colonies. Conversely, in the absence of type IV pili motility, *P. aeruginosa* cannot invade *S. aureus* colonies but rather grows around them, leading to an altered *S. aureus* colony architecture resulting in compact, thicker colonies that have increased biomass compared to coculture with WT *P. aeruginosa*. In addition to these effects on *S. aureus* colony architecture, we also found that coculture with *P. aeruginosa* mutants that lack type IV pili-mediated motility leads to drastically altered physiology through the induction of a significant HQNO-mediated increase in *S. aureus* fermentation. Moreover, type IV pili motility was crucial for modulating the spatial arrangement and competitive dynamics between *P. aeruginosa* and *S. aureus* in conditions that capture essential features of the CF airway environment. Overall, our findings highlight the importance of spatial organization in community-based behaviors and the need for a more thorough understanding of the interplay between polymicrobial communities in the context of infection.

## RESULTS

### Type IV pili are necessary for *P. aeruginosa* invasion into *S. aureus* colonies

We previously reported that *P. aeruginosa* responds to *S. aureus* from a distance by directionally moving towards and surrounding *S. aureus* colonies in a type IV pili motility-mediated manner [38], but how this behavior affects *S. aureus* physiology remained unclear. To test the consequences of *P. aeruginosa* directional motility on *S. aureus*, we first sought to visualize *P. aeruginosa* interactions with *S. aureus* colonies in three dimensions by performing live resonant scanning confocal microscopy of *S. aureus* USA300 JE2 in mono- or coculture with WT *P. aeruginosa* PA14 or a *P. aeruginosa* type IV pili-deficient mutant (Δ*pilA*). Here, *S. aureus* constitutively expressed *sgfp* (pseudocolored orange), while *P. aeruginosa* constitutively expressed *mCherry* (pseudocolored cyan). Bacteria were inoculated between a cover slip and an agarose pad for visualization in the same visual field over time. Imaging was initiated with *S. aureus* and *P. aeruginosa* as single cells, positioned approximately 30 to 50 µm apart to provide sufficient time and distance for *P. aeruginosa* to respond to the presence of *S. aureus*. As previously demonstrated [38], at approximately 5 h, we observed that WT *P. aeruginosa* responds to *S. aureus* by breaking into single cells and moving towards it with type IV pili motility, which eventually leads to *P. aeruginosa* surrounding, invading, and disrupting *S. aureus* cells from the colony (Fig. 1). This invasion is dependent on type IV pili motility, as *P. aeruginosa* Δ*pilA* exhibited significantly decreased invasion compared to WT *P. aeruginosa* (Fig. 1B). While the Δ*pilA* mutant is amotile, it eventually grows against the *S. aureus* colony at later time points (Fig. 1A and C). These data suggest that type IV pili motility is not only necessary for *P. aeruginosa* directional movement towards *S. aureus*, but also aids in the effective invasion of *P. aeruginosa* into *S. aureus* colonies.

**Figure 1.**
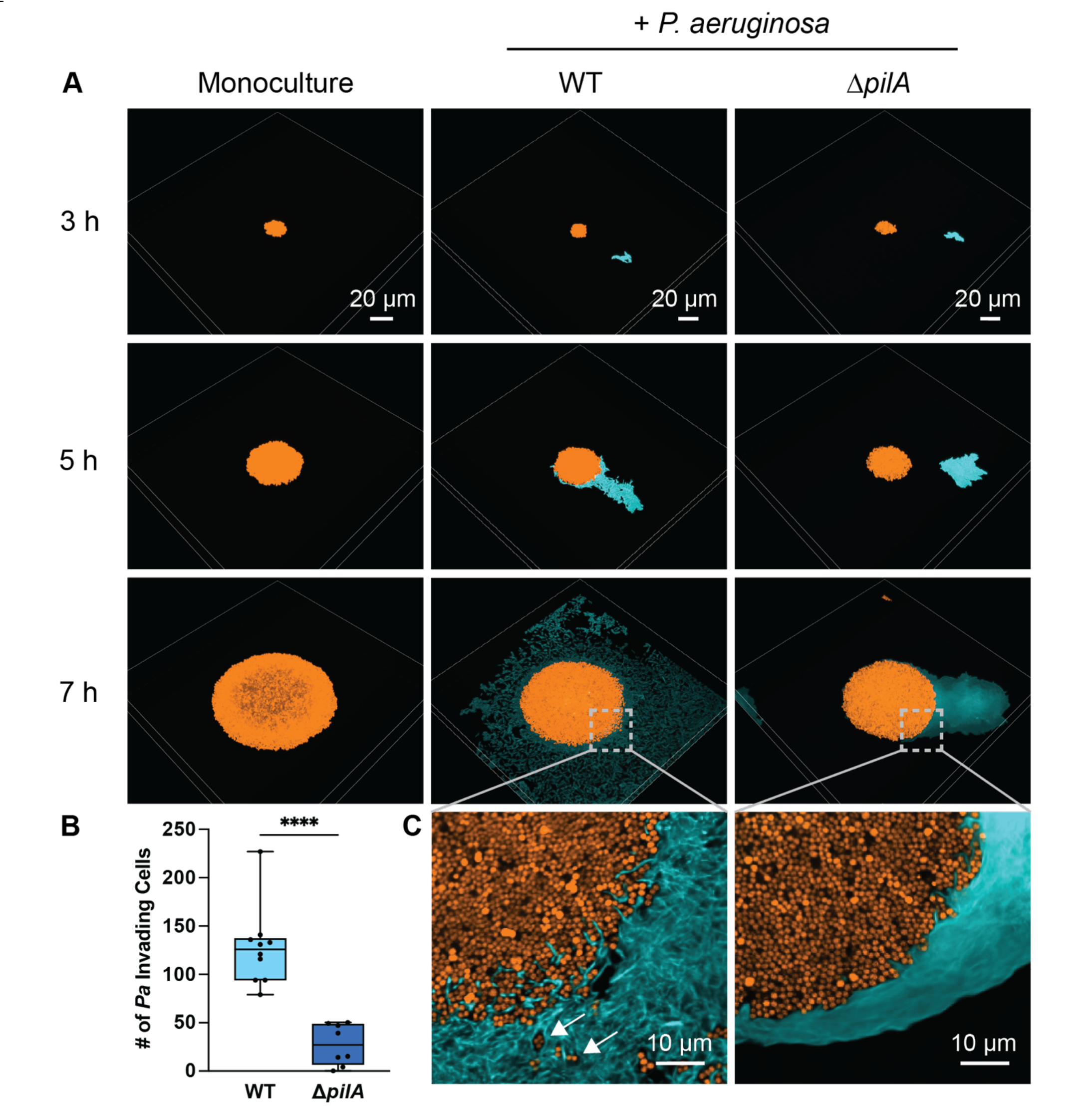
Type IV pili are necessary for *P. aeruginosa* invasion into *S. aureus* colonies. Resonant scanning confocal live imaging of *S. aureus* and *P. aeruginosa*. **A.** Representative micrographs of WT *S. aureus* (pseudocolored orange) in monoculture or in coculture with *P. aeruginosa* (pseudocolored cyan; WT or TFP-deficient mutant Δ*pilA*). **B.** Quantification of *P. aeruginosa* single-cell invasion into *S. aureus* colonies t ∼7 h in mono- or coculture with *P. aeruginosa* (WT or Δ*pilA*). At least four biological replicates with two technical replicates each were analyzed. Each data point represents one technical replicate. Statistical significance was determined by a Mann-Whitney’s test. **** = *P* < 0.0001. **C.** Zoomed micrograph of *S. aureus* colony edge in coculture with *P. aeruginosa* (WT or Δ*pilA*). White arrows indicate dispersed *S. aureus* cells. *S. aureus*: pCM29 P_*sarA*_P1-_*sgfp*_; *P. aeruginosa*: chromosomal P_*A1/04/03-mCherry*_.

### *P. aeruginosa* type IV pili motility-mediated invasion influences *S. aureus* growth and architecture

To investigate how *P. aeruginosa* invasion into *S. aureus* colonies affects *S. aureus* physiology, we imaged *S. aureus* in mono- or coculture with WT or Δ*pilA P. aeruginosa* following ∼24 h of competition. At later time points, visualizing *P. aeruginosa* becomes challenging due to a combination of a decreased fluorescence from the chromosomal P*A1/04/03-mCherry* transcriptional reporter and photobleaching. Nevertheless, we confirmed through phase contrast microscopy that *P. aeruginosa* cells surround *S. aureus* colonies after ∼24 h (Supplementary Fig. 1).

We found that in coculture with WT *P. aeruginosa*, *S. aureus* forms smaller colonies than in monoculture (Fig. 2A), as demonstrated by measuring the area at the base of the *S. aureus* colony (Fig. 2B). Moreover, *P. aeruginosa* type IV pili motility-mediated invasion resulted in *S. aureus* colony edges exhibiting reduced fluorescence, likely caused by dispersed, lysed cells, or a combination thereof (Fig. 2A). In the presence of the non-twitching *P. aeruginosa* strain, the area of *S. aureus* colonies was comparable to that in the presence of WT *P. aeruginosa* (Fig. 2B). However, while the total growth area was similar, *S. aureus* colonies formed during coculture with the Δ*pilA* mutant exhibited less dispersal at the colony edges, possibly due to the absence of *P. aeruginosa* invasion (Fig. 2A; top and middle row). Along with this observation, *S. aureus* colonies appeared thicker and denser than in coculture with WT *P. aeruginosa*. These results suggest that when grown in the presence of the Δ*pilA* mutant, *S. aureus* forms colonies that are more densely packed, possibly due to less dispersal of cells from the colony.

**Figure 2.**
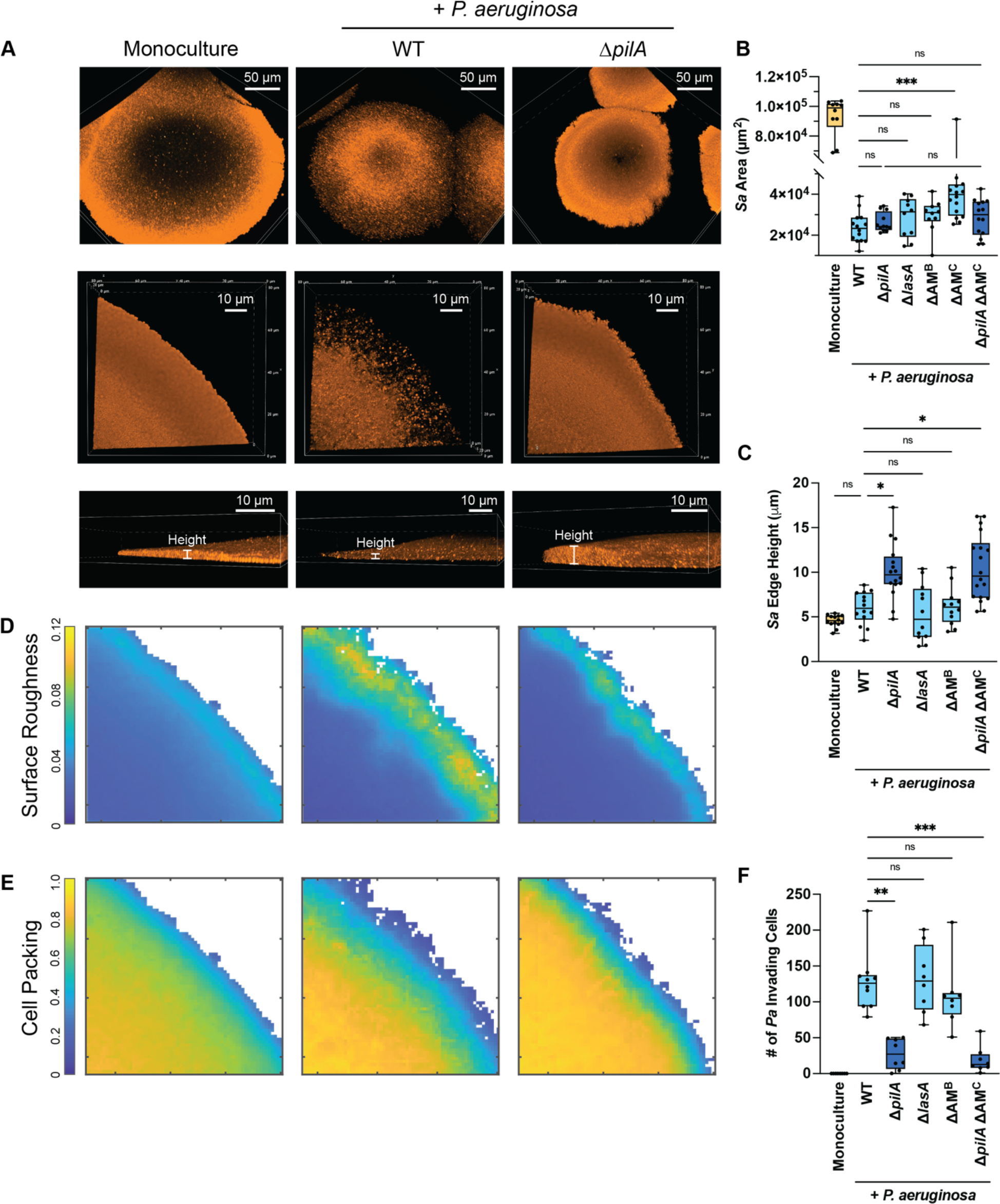
*P. aeruginosa* type IV pili motility-mediated invasion influences the architecture of *S. aureus* colonies independently of *P. aeruginosa*-secreted antimicrobials. Analysis of *S. aureus* colony edge disruption and thickness. **A.** Representative resonant scanning confocal micrographs of the whole colony (top row) or Galvano scanner colony edge micrographs of WT *S. aureus* (orange) in monoculture or coculture with *P. aeruginosa* (not shown; WT or Δ*pilA*) t ∼24 h shown from the top (middle row) of the colony or the side (bottom row). The micrographs in **A** (bottom row) show the colonies on the Z-plane and demonstrate how the height was quantified. **B & C.** Quantification of *S. aureus* whole colony area at t ∼24 h (**B**), or height at the edge of *S. aureus* (*Sa*) colony (µm) at t ∼24 h (**C**) in monoculture or coculture with *P. aeruginosa* (WT, Δ*pilA*, ΔAM^B^ (bacteriostatic antimicrobials; HQNO, pyoverdine, and pyochelin), Δ*lasA*, ΔAM^C^ (complete antimicrobials; HQNO, pyoverdine, pyochelin, and LasA), or Δ*pilA* ΔAM^C^). **D & E.** Representative BiofilmQ heatmaps of local surface roughness (**D**) and cell packing analysis (**E**) at *S. aureus* colony edge in mono- or coculture with *P. aeruginosa* (WT or Δ*pilA*). Three biological replicates with two technical replicates each were analyzed in D & E. **F.** The number of invading *P. aeruginosa* (*Pa*) single cells found inside *S. aureus* was quantified at t ∼7 h in mono- or coculture with the *P. aeruginosa* strains described above. At least four biological replicates with two technical replicates each were analyzed per condition in B, C, and F. Each data point represents one technical replicate. Statistical significance was determined by Kruskal-Wallis followed by Dunn’s Multiple Comparisons Test. n.s. = not significant; * = *P* < 0.05; ** = *P* < 0.01; *** = *P* < 0.001.

To test this hypothesis, we sought to visualize and quantify *S. aureus* colony architecture in more detail. Since colony thickness and density were more distinct on the edge of the colonies, images were also acquired with higher magnification and increased spatial resolution using galvanometric point-scanning confocal microscopy at a single final time point (Fig. 2A; middle row). We then measured the height and density at the edge of *S. aureus* colonies at 15 µm from the edge using the Z-plane as illustrated in Fig. 2A (bottom row) and quantified in Fig. 2C. As expected, the height at the edge of the colony was significantly higher in coculture with Δ*pilA P. aeruginosa* than in mono- or coculture with WT *P. aeruginosa* (Fig. 2C). To obtain a quantitative analysis of the colony density, we measured both cell packing and colony surface roughness using the microscopy image analysis software BiofilmQ [41]. These parameters (cell packing and surface roughness) quantify density by measuring the amount of surface or volume within a specified area. In BiofilmQ, *S. aureus* colony edges were separated from the background by segmentation and dissection onto a 3D grid, with each cubic grid unit measuring 0.72 µm on each side. The neighborhood surface roughness and cell packing were then calculated by determining the biovolume fraction and surface height variance of *S. aureus* for each grid cube within 4 and 6 µm, respectively. Representative heatmaps in Fig. 2D and 2E provide visualization of the quantified data in two-dimensions, using color coding to represent local surface roughness and cell packing. We observed that *S. aureus* colonies in monoculture have low surface roughness and are uniformly packed together (Fig. 2D and E; first column). In contrast, when WT *P. aeruginosa* is present, *S. aureus* edges exhibit increased surface roughness and decreased cell packing (Fig. 2D and E; middle column), which suggests reduced colony density caused by WT *P. aeruginosa*. We also assessed *S. aureus* cell packing following coculture with the Δ*pilA* mutant (i.e., in the absence of *P. aeruginosa* invasion and colony disruption). This analysis revealed increased cell packing and reduced colony surface roughness compared to WT *P. aeruginosa* coculture, much like *S. aureus* monoculture colonies (Fig. 2D and E). Thus, while the area at the base of the colony is similar in the presence of WT or Δ*pilA*, the colonies are more densely packed when *P. aeruginosa* lacks type IV pili motility. Altogether, these observations reveal the crucial role of *P. aeruginosa* type IV pili motility in altering *S. aureus* architecture. Without type IV pili motility, *P. aeruginosa* exhibits reduced invasion and disruption of *S. aureus* colonies; instead, it grows alongside them, which results in increased compaction and changes in the structure of *S. aureus* colonies.

### *P. aeruginosa* secretes antimicrobials that affect *S. aureus* growth but do not influence *S. aureus* colony architecture

Next, we wondered how invasion changes *S. aureus* colony architecture and enhances competition. Invasion of *S. aureus* colonies by *P. aeruginosa* results in reduced *S. aureus* colony density (Fig. 2); one potential explanation for this is that invasion increases the local concentration of *P. aeruginosa* antimicrobials within *S. aureus* colonies. It is also possible that these anti-staphylococcal factors actually aid in *P. aeruginosa* invasion into *S. aureus* colonies. If the former is correct, we would expect *S. aureus* colonies grown in the presence of the *P. aeruginosa* type IV pili mutant to be protected from *P. aeruginosa* antimicrobials. *P. aeruginosa* secretes a large number of antimicrobial compounds known to inhibit or lyse *S. aureus*, including 2-heptyl-4-hydroxyquinoline *N*-oxide (HQNO), a respiratory toxin that inhibits the *S. aureus* electron transport chain [26, 32], the siderophores pyoverdine and pyochelin, which aid in iron scavenging [25, 28], and a potent anti-staphylococcal protease, staphylolysin or LasA, which lyses *S. aureus* by cleaving the peptidoglycan pentaglycine cross-links [27].

Since the *P. aeruginosa* type IV pili motility mutant was found to have reduced invasion and disruption of *S. aureus* exterior structure compared to WT *P. aeruginosa*, we first sought to determine if *P. aeruginosa* exoproducts affected this phenotype, and tested whether the Δ*pilA* mutant produces similar levels of secreted antimicrobials as the WT. To address this, the cell-free supernatant from the Δ*pilA* mutant and WT *P. aeruginosa* was added to *S. aureus* to examine its growth and lysis over time. No differences were observed in either *S. aureus* lysis or growth rate when exposed to the supernatant from WT or Δ*pilA P. aeruginosa*, confirming that the Δ*pilA* strain produces antimicrobials at the same level as WT (Supplementary Fig. 2). For this lysis assay, the supernatant from the *P. aeruginosa* Δ*lasA* mutant served as a control to confirm that staphylolysin is the main driver of *S. aureus* lysis under these conditions.

To test the hypothesis that invasion by motile *P. aeruginosa* cells enhances competition by increasing diffusion and thus local concentration of *P. aeruginosa* antimicrobials within *S. aureus* colonies, we next examined the colony growth dynamics of *S. aureus* in the presence of *P. aeruginosa* strains carrying clean deletions of genes encoding secreted products. These include a strain that lacked the production of both HQNO and siderophores (Δ*pqsL* Δ*pvdA* Δ*pchE*), which we refer to as “ΔAM^B^” (antimicrobials^Bacteriostatic^) in Fig. 2B, 2C, and 2F, and a *P. aeruginosa* strain unable to produce the bacteriolytic enzyme staphylolysin (Δ*lasA*). The interactions between *S. aureus* and these antimicrobial-deficient *P. aeruginosa* strains were assessed by live-imaging as described for Figure 1, and *P. aeruginosa* invasion and *S. aureus* colony height (as a proxy for biomass) were quantified. No detectable differences were observed between coculture with WT *P. aeruginosa* and the antimicrobial mutants for either the *S. aureus* colony edge height (Fig. 2C) or the number of invading cells (Fig. 2F), which suggests that these antimicrobials do not play a role in *P. aeruginosa* invasion or in the increased *S. aureus* colony height observed in the presence of the Δ*pilA* mutant strain. Yet, it is known that these antimicrobials can impact *S. aureus* growth *in vitro* [7, 9, 11, 12, 31, 33, 34, 42]. Therefore, we measured the area at the base of the colony of *S. aureus* to examine if these antimicrobials influence *S. aureus* growth under the current experimental conditions. *S. aureus* colony area did not significantly increase when cocultured with the *P. aeruginosa* strains that lacked either the bacteriostatic antimicrobials or staphylolysin alone, compared to WT *P. aeruginosa* (Fig. 2B). However, colony area did increase upon deletion of all four antimicrobials (HQNO, both siderophores, and staphylolysin) simultaneously from *P. aeruginosa*, which we refer to as “ΔAM^C^” (AM^Complete^ mutant, Fig. 2B). These data suggest that while these antimicrobials do not influence *S. aureus* colony architecture, their combinatorial effect does alter *S. aureus* growth and colony area under these conditions.

To determine if the changes in *S. aureus* colony edge height and *P. aeruginosa* invasion are driven by motility alone or a combined effect of motility and antimicrobials, we deleted *pilA* in the ΔAM^C^ mutant, generating Δ*pqsL* Δ*pvdA* Δ*pchE* Δ*lasA* Δ*pilA* (Δ*pilA* ΔAM^C^) in Fig. 2B, 2C, and 2F. *S. aureus* colony height and *P. aeruginosa* invasion in the presence of Δ*pilA* ΔAM^C^ phenocopied that of the Δ*pilA* mutant (Fig. 2C and F), suggesting that type IV pilus motility plays a prominent role in driving these phenotypes. Furthermore, when comparing the effect of deletion of *pilA* in the WT background to the complete AM background, a significant influence of antimicrobials on colony area is only observed when type IV pili are functional, supporting the hypothesis that motility may enhance AM action against *S. aureus* under these conditions (Fig. 2B).

Overall, these data suggest that the change in *S. aureus* colony architecture becoming thicker and denser is exclusively mediated by the absence of *P. aeruginosa* type IV pili-mediated colony invasion and that the main *P. aeruginosa* anti-staphylococcal factors do not substantially influence this observation. Furthermore, *P. aeruginosa* type IV pili motility may enhance antimicrobial access into the colony to fully affect *S. aureus* growth, revealing the important role *P. aeruginosa* motility plays in antagonistic interactions against *S. aureus*.

### Increased cell packing enhances HQNO-mediated *S. aureus* fermentation

Although *P. aeruginosa* antimicrobials did not influence *S. aureus* structure, we next sought to explore how *S. aureus* change in colony morphology affects its physiological response to HQNO by utilizing a fluorescent reporter system. HQNO poisons the *S. aureus* respiratory chain, forcing a shift to fermentative metabolism [7], and so the extent of *S. aureus* fermentation can be used as a proxy for HQNO activity. Fermentation in turn can be measured via a fluorescent transcriptional fusion to the promoter of the gene encoding lactate dehydrogenase (P*ldh1*-*sgfp*) [43]. If HQNO penetrates densely packed *S. aureus* colonies, we would expect to see an increase in fluorescence compared to coculture with a *P. aeruginosa* mutant that lacks HQNO production. To test this prediction, we live-imaged *S. aureus* in mono- or coculture with *P. aeruginosa* and quantified the mean fluorescence intensity (MFI; fluorescence / colony volume) per *S. aureus* colony over an 18-hour period (Fig. 3). *S. aureus* fermentation in coculture with WT *P. aeruginosa* began to increase at approximately 12 h. P*ldh1*-*sgfp* fluorescence did not increase when grown in the presence of Δ*pqsL*, confirming prior reports that HQNO increases *ldh* expression [7]. To test if fermentation increases in the absence of *P. aeruginosa* invasion, *S. aureus* P*ldh1*-*sgfp* expression was quantified in coculture with *P. aeruginosa* Δ*pilA*. Notably, we observed a sharp increase in fermentation within the densely packed colonies produced by coculture with the non-motile *P. aeruginosa*, as shown in Fig. 3A and quantified in Fig. 3B and 3C. One interpretation of these data is that HQNO concentrates within densely packed colonies, inducing a more dramatic change in *S. aureus* physiology. An alternative explanation is that the increased P*ldh1*-*sgfp* signal results from increased density, independent of HQNO, potentially due to oxygen restriction within the colony. To differentiate these possibilities, fermentation was measured in the presence of a double Δ*pqsL* Δ*pilA* mutant. As seen with motile *P. aeruginosa* lacking *pqsL*, the double mutant does not induce *S. aureus* fermentation over time (Fig. 3B), suggesting that the increase in fermentation observed is mediated by HQNO. Importantly, the phenotypes of both the Δ*pilA* and Δ*pqsL* mutants could be genetically complemented when their respective genes were expressed in *cis* (for Δ*pilA*) or *trans* (for Δ*pqsL*) under control of an inducible promoter (Supplementary Fig. 3). These findings show that HQNO can likely diffuse into *S. aureus* colonies independently of *P. aeruginosa* invasion and plays a crucial role in mediating interspecies interactions by pushing *S. aureus* towards fermentation.

**Figure 3.**
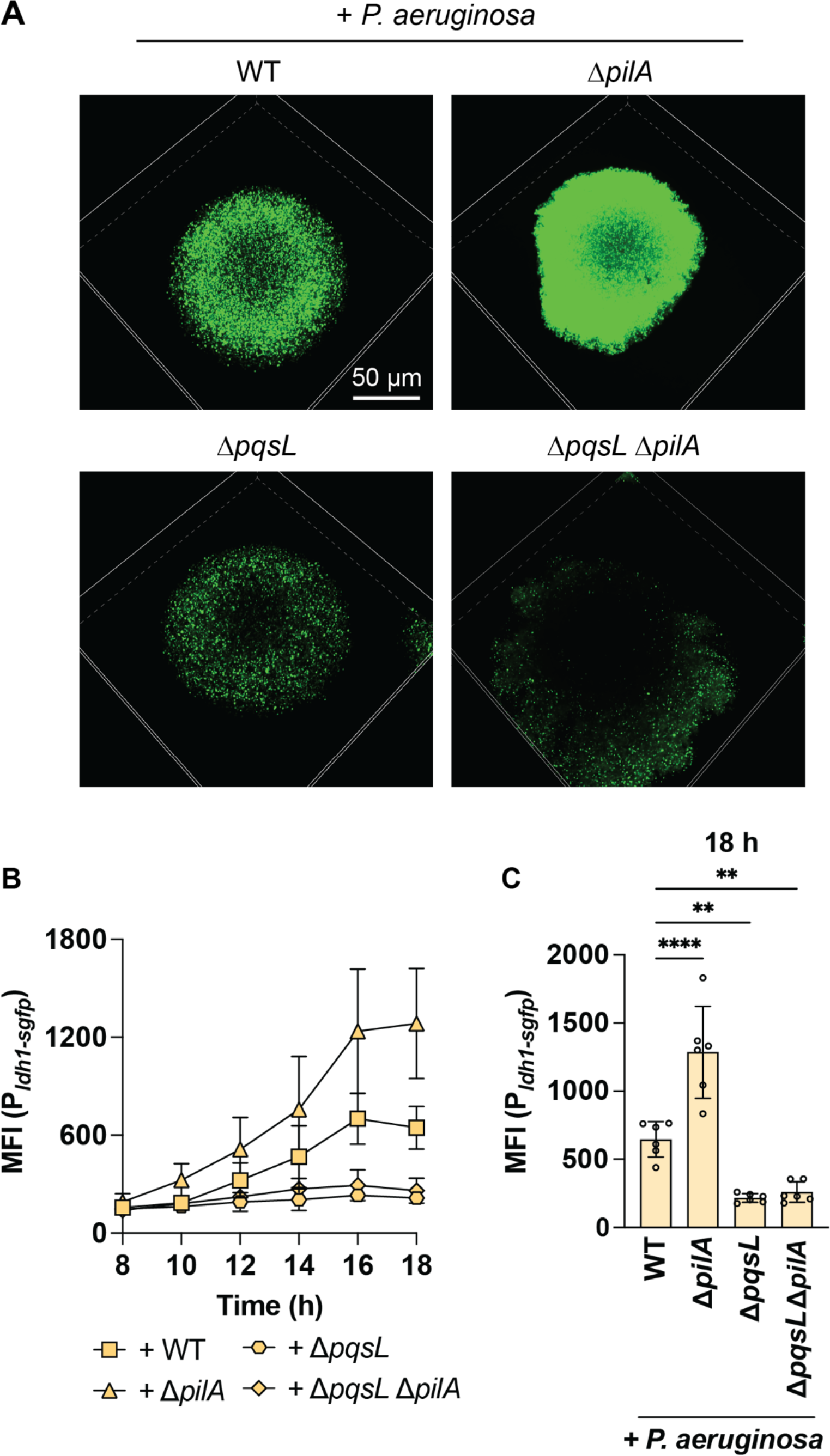
Increased *S. aureus* cell packing enhances HQNO-mediated *S. aureus* fermentation. *S. aureus* lactate fermentation (P_*ldh1-sgfp*_) was measured in the presence of the indicated *P. aeruginosa* strains. **A.** Representative resonant scanning confocal micrographs of *S. aureus* fermentation in coculture with *P. aeruginosa* (WT, Δ*pilA*, Δ*pqsL*, or Δ*pqsL* Δ*pilA*) t = 18 h, *S. aureus* channel only. **B.** Mean fluorescence intensity (MFI, fluorescence / colony volume) was quantified over time. **C.** MFI at 18 h. Data represent the mean and standard deviation from three biological replicates with two technical replicates per condition. Statistical analyses were performed at 18 h using one-way ANOVA followed by Dunnett’s Multiple Comparisons Test comparing each condition to +WT *P. aeruginosa*. ** = *P* < 0.01; **** = *P* < 0.0001.

### *P. aeruginosa* type IV pili motility is necessary for competition against *S. aureus* in conditions that mimic cystic fibrosis lung secretions

While experiments thus far reveal a role for type IV pili motility in competition against *S. aureus*, the imaging conditions utilized constrain cells to the surface between a coverslip and an agarose pad. While useful for high spatial and temporal resolution of interspecies interactions, this approach does not accurately reflect other attributes of the CF airway infection environment. Thus, we sought to determine if *P. aeruginosa* type IV pili motility is an important driver of interspecies interactions when cocultured under conditions that mimic CF lung secretions. To explore this question, *P. aeruginosa* and *S. aureus* were cocultured in Artificial Sputum Media (ASM), a modified version of SCFM2 [44]. ASM captures some of the essential features of the CF environment, like constraints on movement and diffusion, that shaken liquid culture methods do not, and similar recipes have been shown to recapitulate approximately 86% of *P. aeruginosa* gene expression in human expectorated CF sputum, outperforming both laboratory media and the acute mouse pneumonia model of infection [45, 46].

ASM was inoculated with *S. aureus* and *P. aeruginosa* (WT or Δ*pilA*) at a 1:1 ratio, grown statically at 37°C for 22 h, and imaged with resonant scanning confocal microscopy to visualize their spatial organization. The initial inoculum and end time point cultures were plated for colony-forming units (CFUs) to assess bacterial viability. In the presence of WT motile *P. aeruginosa*, *S. aureus* was suppressed relative to its monoculture condition: very few *S. aureus* cells could be observed by microscopy (Fig. 4A), or by viability plating (less than limit of detection) (Fig. 4B), compared to approximately 10^8^ CFUs/well recovered for *S. aureus* in monoculture. However, when *S. aureus* was grown in the presence of the *P. aeruginosa* type IV pili-deficient strain, Δ*pilA*, a 100 - 10,000-fold increase in *S. aureus* cells was recovered in comparison to coculture with WT *P. aeruginosa* (Fig. 4B). Overall, this suggests that type IV pili motility is necessary for effective competition with *S. aureus* under these conditions. Since type IV pili are also necessary for *P. aeruginosa* biofilm formation and attachment to surfaces [47–49], it is formally possible that differences in biofilm formation between the WT and Δ*pilA* strains could influence competition with *S. aureus*. However, *P. aeruginosa* biofilm formation and spatial organization were similar in appearance between these strains in coculture with *S. aureus* in ASM (Fig. 4A). While the CFUs/well recovered for Δ*pilA* were significantly lower than WT *P. aeruginosa* in mono- or coculture with *S. aureus* (Fig. 4C), the difference is modest (∼95% of WT) and thus not expected to account for the increase in *S. aureus* survival. Next, we tested if the mere presence of type IV pili has a role in competition or if type IV pili motility is required. To differentiate between these two outcomes, a hyperpiliated, non-twitching *P. aeruginosa* mutant (Δ*pilT*) was cocultured with *S. aureus*. This mutant lacks the main retraction ATPase of the type IV pili machinery, PilT, and is well-documented to ineffectively retract extended pili [50]. *S. aureus* survival in the presence of *P. aeruginosa* Δ*pilT* phenocopied survival in the presence of Δ*pilA* (Fig. 4A and B), suggesting that *P. aeruginosa* functional type IV pili are necessary for competitive interactions with *S. aureus* in CF sputum. Collectively, these data demonstrate that under CF-relevant conditions, *P. aeruginosa* type IV pili motility aids in interspecies competition.

**Figure 4.**
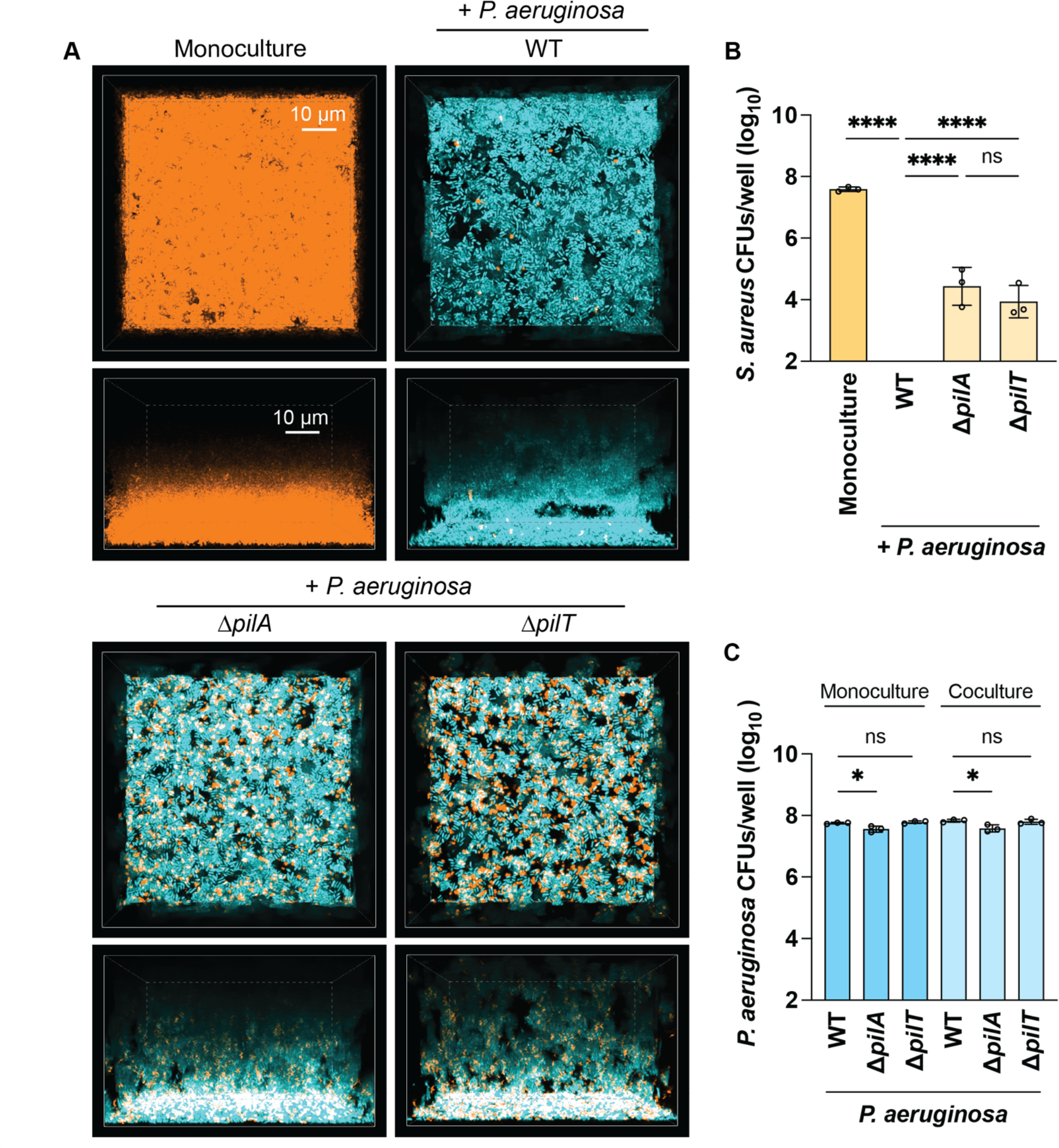
*P. aeruginosa* type IV pili motility is necessary for competition against *S. aureus* in artificial sputum media. Resonant scanning confocal imaging of *S. aureus* and *P. aeruginosa* under static conditions in artificial sputum media, with CFU quantification. **A.** Representative images of resonant scanning confocal micrographs of WT *S. aureus* (pseudocolored orange) in monoculture or in coculture with *P. aeruginosa* (pseudocolored cyan; WT, Δ*pilA*, or Δ*pilT*) t ∼22 h. *S. aureus* (**B**) or *P. aeruginosa* (**C**) CFU quantification in monoculture or in coculture with *P. aeruginosa* (WT, Δ*pilA*, or Δ*pilT*) (**B**), or in mono- or coculture with *S. aureus* (**C**) at t = 24 h. The CFUs/well in Y-axes are portrayed as log10 transformed. Three biological replicates with one technical replicate each were analyzed, and the mean and standard deviation are shown. Each data point represents one biological replicate. Statistical significance was determined by one-way ANOVA followed by Dunnett’s Multiple Comparisons Test. n.s. = not significant; * = *P* < 0.05; **** = *P* < 0.0001.

### *P. aeruginosa* type IV pili motility is necessary for disruption and competition against pre-formed *S. aureus* biofilms

Next, we asked if *P. aeruginosa* type IV pili motility is necessary for competition against pre-formed *S. aureus* biofilms in ASM. *S. aureus* was grown statically at 37°C in ASM for approximately 5 h. Subsequently, *P. aeruginosa* WT, Δ*pilA*, or Δ*pilT* were added to *S. aureus* and allowed to grow for an additional 24 h in coculture. Then, the bacteria were imaged and plated for viability. Remarkably, we found that WT *P. aeruginosa* can invade and disrupt pre-formed *S. aureus* biofilms, as depicted in Fig. 5A (Z plane) where a layer of the *S. aureus* biofilms can be seen detached from the surface and blanketed by *P. aeruginosa* cells. *S. aureus* disruption was dependent on *P. aeruginosa* type IV pili motility, as WT *P. aeruginosa* disrupted *S. aureus* pre-formed biofilms significantly more than the non-twitching *P. aeruginosa* mutants (Supplementary Fig. 4). Notably, S*. aureus* and *P. aeruginosa* Δ*pilA* remained segregated into monoculture aggregates, whilst cells were well-mixed during coculture with motile *P. aeruginosa* (Fig. 5A, inset). These observations were consistent with the previous data acquired with the experimental conditions under agarose pads. We also observed that a higher number of WT *P. aeruginosa* colonized the surface of the coverslip compared to the Δ*pilA* or Δ*pilT* mutants in the presence of *S. aureus* (Fig. 5A and Supplementary Fig. 4). These data suggest that type IV pili motility is necessary for *P. aeruginosa* cells to invade from the top of *S. aureus* biofilms and traverse through to access the coverslip, potentially disrupting and detaching the biofilms in the process. This results in a significant reduction in *S. aureus* viability (10 - 15-fold) when comparing *S. aureus* coculture with WT vs Δ*pilA* and Δ*pilT* mutants (Fig. 5B). Importantly, no viability differences were observed between the WT *P. aeruginosa* and either of the TFP mutants (Fig. 5C). Altogether, these observations suggest that type IV pili motility enhances *P. aeruginosa* competitive fitness, allowing it to disrupt and potentially render *S. aureus* cells more vulnerable to *P. aeruginosa* antimicrobials.

**Figure 5.**
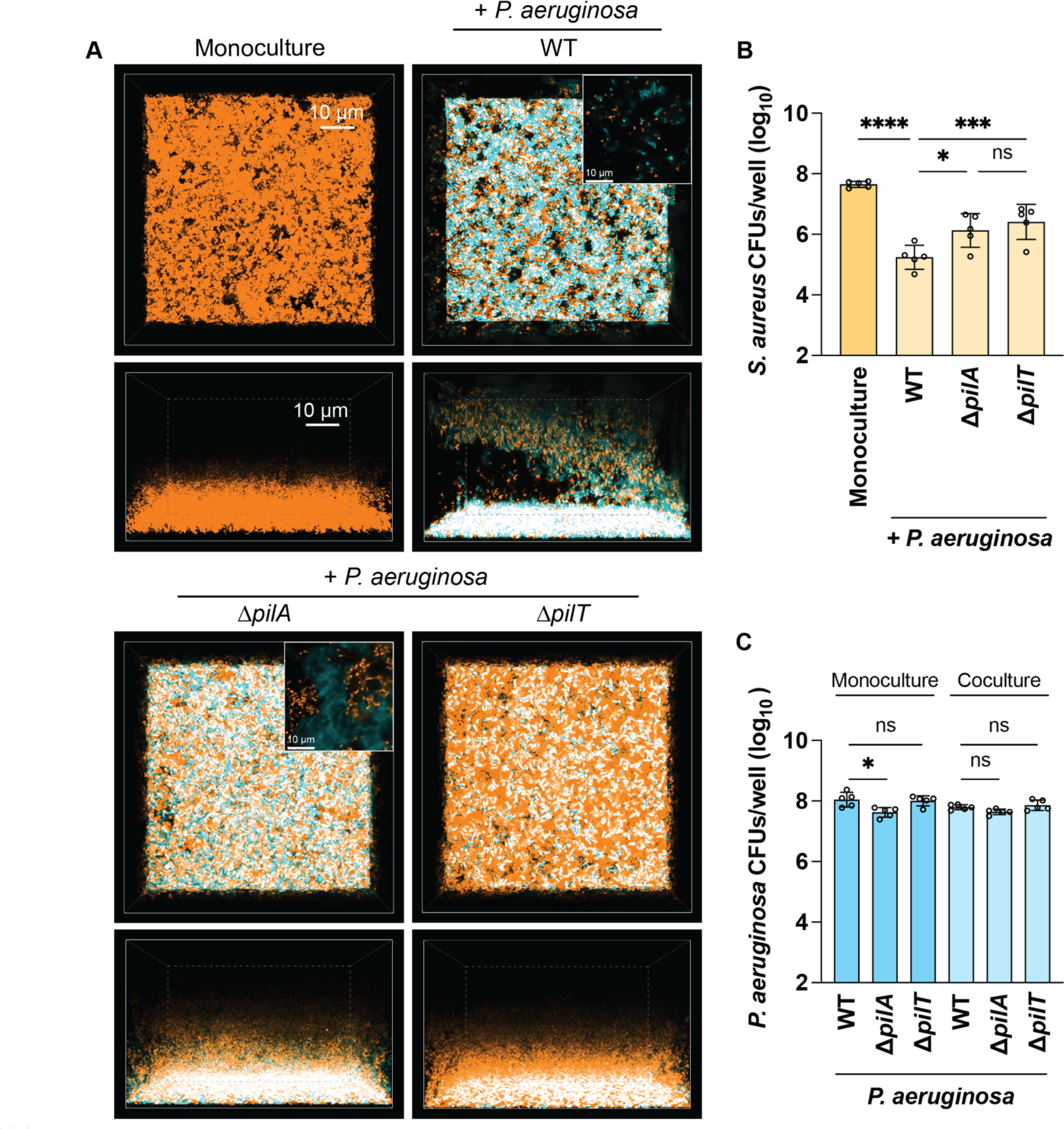
*P. aeruginosa* type IV pili motility is necessary for disruption and competition against pre-formed *S. aureus* biofilms. Resonant scanning confocal imaging of *S. aureus* and *P. aeruginosa* under static conditions in artificial sputum media, with CFU quantification at late time points. *P. aeruginosa* was added to *S. aureus* pre-formed biofilms at 5 h. **A.** Representative images of resonant scanning confocal micrographs of WT *S. aureus* (pseudocolored orange) in monoculture or in coculture with *P. aeruginosa* (pseudocolored cyan; WT, Δ*pilA*, or Δ*pilT*) t ∼26 h. The insets show ∼3X zoomed images at 30 μm from the base of the coverslip. *S. aureus* (**B**) or *P. aeruginosa* (**C**) CFU quantification in mono- or in coculture with *P. aeruginosa* (WT, Δ*pilA*, or Δ*pilT*) (**B**), or in mono- or coculture with *S. aureus* (**C**) at t ∼29 h. The CFUs/well in Y-axes are portrayed as log10 transformed. Five biological replicates with one technical replicate each were analyzed, and the mean and standard deviation are shown. Statistical significance was determined by one-way ANOVA followed by Dunnett’s Multiple Comparisons Test. n.s. = not significant; * = *P* < 0.05; *** = *P* < 0.001; **** = *P* < 0.0001.

## DISCUSSION

Growing data support the hypothesis that spatial organization is an essential determinant in shaping microbial communities and can further dictate important community-based behaviors [4, 5, 35, 51–57]. In this study, we found that *P. aeruginosa* motility plays a vital role in shaping the biogeography in polymicrobial environments with *S. aureus*. By influencing spatial aggregation, *P. aeruginosa* type IV pili motility ultimately determines *S. aureus* physiology and survival.

While antimicrobials secreted by *P. aeruginosa* have been extensively documented to influence *S. aureus* growth and survival [7–12], *P. aeruginosa* motility in interspecies competition has only begun to be explored. We recently reported that *P. aeruginosa* can sense *S. aureus* secreted peptides, phenol soluble modulins (PSMs), from a distance via the PilJ methyl-accepting chemoreceptor [40]. Consequently, it employs twitching motility to chemotax towards *S. aureus* colonies or PSMs alone [38, 39]. In addition to the chemosensory response, *S. aureus* PSMs also trigger a “competition sensing” response whereby *P. aeruginosa* upregulates type VI secretion system and pyoverdine biosynthesis pathways [39]. Similarly, *P. aeruginosa* has been reported to utilize type IV pili-mediated motility to perform “suicidal chemotaxis” toward antibiotics [58]. The upregulation of these common interbacterial competition pathways supports a model where *P. aeruginosa* senses potential danger in the environment and responds with directional twitching, while simultaneously activating defense mechanisms to combat the “enemy”. Additional evidence that supports this model has been reported wherein *P. aeruginosa* upregulates antimicrobial production upon sensing *N*-acetylglucosamine alone or shed from Gram-positive bacteria [59].

Our single-cell level temporal analysis also revealed that *P. aeruginosa* type IV pili motility is necessary for invading and disrupting *S. aureus* colonies (Fig. 1). Interestingly, loss of invasion leads *P. aeruginosa* to grow adjacent to *S. aureus* colonies and potentially acting as a “wall” that may prevent expansion of the *S. aureus* colonies, which become thicker and denser (Fig. 2). While *P. aeruginosa* anti-staphylococcal factors did not mediate invasion into *S. aureus* colonies, they did influence *S. aureus* growth as *S. aureus* formed larger colonies in the absence of *P. aeruginosa* antimicrobials HQNO, pyoverdine, pyochelin, and staphylolysin (Fig. 2B). It was not surprising that deletion of all four antimicrobials was required to observe enhanced *S. aureus* growth, since *P. aeruginosa* effects on *S. aureus* have been well-documented to depend on multiple secreted antimicrobials simultaneously such as HQNO, pyoverdine, and pyochelin [7, 9, 12]. However, most of these previous studies were performed with *P. aeruginosa* cell-free supernatant, and not with live *P. aeruginosa* present. Imaging *P. aeruginosa* and *S. aureus* in coculture at the single-cell level has allowed us to visualize the importance of *P. aeruginosa* motility in their interactions, and therefore, start to build a model whereby twitching motility aids in competition by disrupting *S. aureus* single cells away from the colony, leaving them exposed and potentially more vulnerable to *P. aeruginosa*-secreted factors (Fig. 6). Therefore, when *P. aeruginosa* cannot move, we hypothesize that *S. aureus* cells remain protected within the colony and resist infiltration of *P. aeruginosa* antimicrobials. Altogether, these findings provide additional support of how type IV pili motility can either enhance competition or foster coexistence with *S. aureus*.

**Figure 6.**
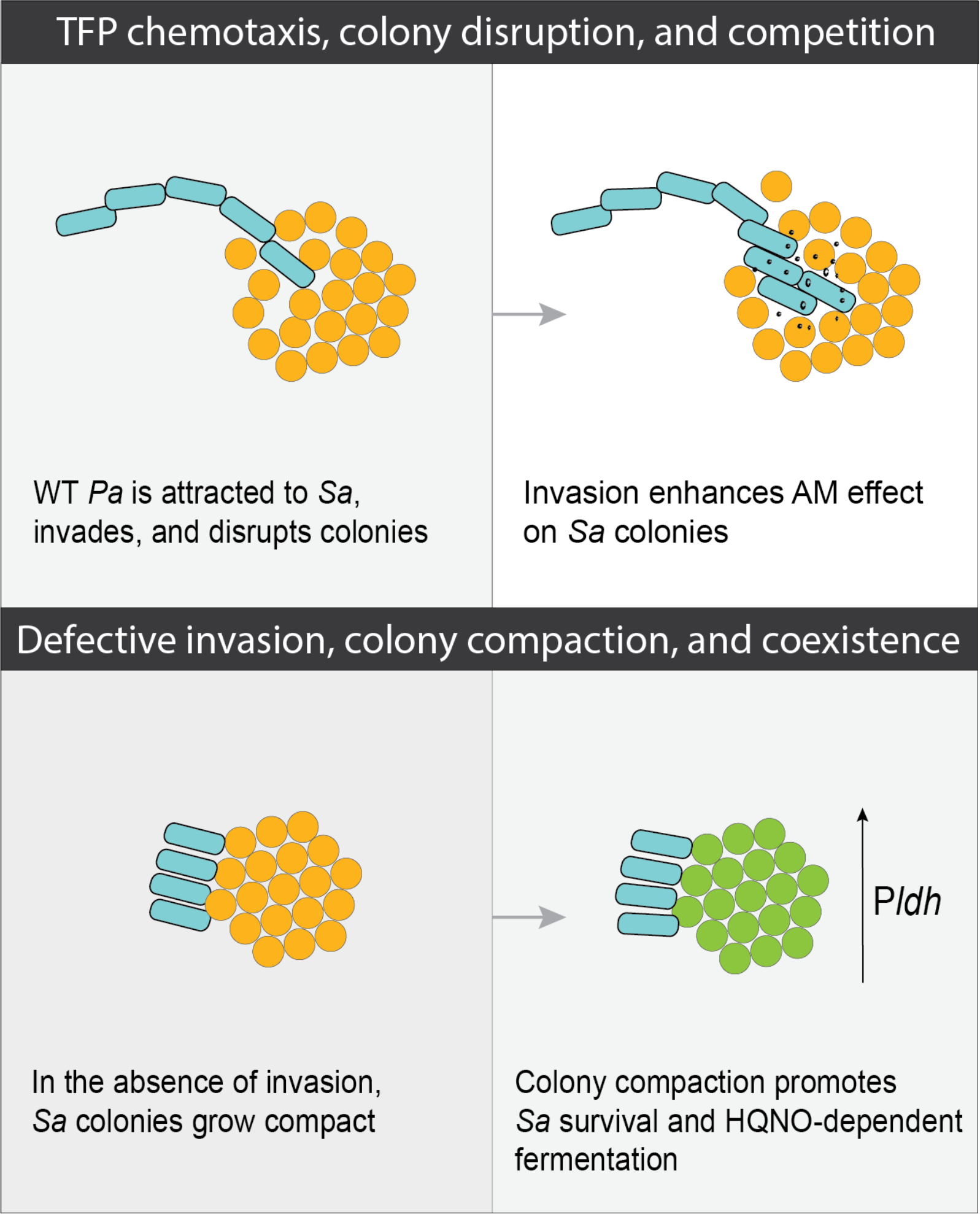
Model for motility-driven interspecies competition. We propose that *P. aeruginosa* type IV pili motility-mediated attraction towards, invasion, and disruption of *S. aureus* colonies promotes the diffusion of antimicrobials (AM) to maximize interspecies competition. Lack of motility affects *S. aureus* spatial organization and physiology in a manner that promotes coexistence.

*S. aureus* colonies with a different morphology arise as a consequence of the limited invasion of *P. aeruginosa* Δ*pilA* and show a striking change in physiology by increasing lactic acid fermentation (P*ldh1*-*sgfp*) compared to those cocultured with WT *P. aeruginosa* (Fig. 3). While we initially hypothesized that *S. aureus* cells remained protected from *P. aeruginosa* antimicrobials in the absence of *P. aeruginosa* invasion, these data suggest that HQNO can diffuse into *S. aureus* colonies without relying on *P. aeruginosa* type IV pili motility-mediated invasion. Therefore, while *P. aeruginosa* Δ*pilA* cannot invade and disrupt, the antimicrobials it secretes can still alter *S. aureus* growth and physiology. Nevertheless, deletion of antimicrobial production in the Δ*pilA* mutant background does not significantly improve *S. aureus* survival, as it does in the WT background. This observation provides evidence for the role of type IV pili motility invasion and disruption in allowing the access of these antimicrobials into *S. aureus* colonies to have a greater impact on its growth (Fig. 2B). Therefore, we hypothesize that in the absence of TFP-mediated invasion and disruption, antimicrobials with higher molecular weight, such as staphylolysin (36 kDa), are precluded from freely diffusing into *S. aureus* colonies, while smaller compounds like HQNO (0.259 kDa) and possibly siderophores, can diffuse and concentrate within *S. aureus*, eliciting physiological changes that could pose greater challenges for the effective treatment of infections.

Importantly, the relevance of *P. aeruginosa* type IV pili motility in mediating interspecies competition and spatial aggregation was also highlighted under conditions that mimic the nutritional and viscoelastic properties of CF airways. Of note, *P. aeruginosa* was capable of detaching preformed *S. aureus* biofilms and significantly reducing *S. aureus* viability in a motility-dependent manner when grown in a modified synthetic CF sputum medium. These results emphasize the importance of motility in this polymicrobial environment that mimics the conditions in CF airways.

The mucoid *P. aeruginosa* phenotype, which is a common adaptation that *P. aeruginosa* exhibits during CF infections, is associated with decreased competition against *S. aureus* due to the reduced production of anti-staphylococcal factors by these strains [42]. Additional studies have demonstrated that another commonly reported *P. aeruginosa* adaptation linked to chronic CF infections is reduced twitching motility [60]. Interestingly, both *P. aeruginosa* mucoid and reduced twitching phenotypes were identified as the best phenotypic predictors of future pulmonary exacerbations in infected children with CF [60]. Our studies revealed that impairing twitching motility hinders *P. aeruginosa* competitiveness and promotes coexistence with *S. aureus* under CF-relevant conditions. This may be a contributing explanation for why *P. aeruginosa* and *S. aureus* are still found in high numbers in people with CF [14], given that multiple adaptations acquired by chronic *P. aeruginosa* isolates can coexist better with *S. aureus*.

It has been extensively discussed whether *P. aeruginosa* and *S. aureus* encounter each other in the CF lungs, and if they compete or coexist in this setting [4, 35, 42, 51]. CF airways are indeed a complex environment with multiple, distinct niches within; therefore, we predict that *P. aeruginosa* and *S. aureus* can coexist in some areas, while there might be other areas where *P. aeruginosa* could outcompete *S. aureus*. There could be spaces where they are found well-mixed, and others where they are more spatially segregated but still influence each other through the diffusion of secreted factors. As evidenced by both this study and others, it is becoming clear that interspecies competition - or coexistence - can greatly depend on *P. aeruginosa* genotype and phenotype, which can vary in different areas of the lung. Our *in vitro* data show how spatial organization can determine the outcome of microbe-microbe interactions and also help inform the potential interactions of these bacteria during infection. However, the CF airways are complex and involve not only other microorganisms but also host factors that need to be considered. Therefore, future experiments must be performed *in vivo* and *ex vivo* to map the biogeography of these communities and further elucidate these interspecies interactions.

Overall, our study reveals how *P. aeruginosa* type IV pili motility aids in the disruption of *S. aureus* colonies and biofilms, which potentiates the effect of *P. aeruginosa*-secreted anti-staphylococcal factors on *S. aureus* cells. Ultimately, *P. aeruginosa* motility plays a crucial and previously unexplored role in determining *S. aureus* outcome.

## MATERIALS AND METHODS

### Bacterial strains and growth conditions

A list of all the strains used in this study can be found in Table S1 in the supplemental material. For microscopy assays, *P. aeruginosa* (PA14) and *S. aureus* (USA300 JE2) were grown in the minimal medium M8T (M8 salts supplemented with 0.2% glucose, 1.2% tryptone, and 1 mM MgSO4). *S. aureus* fluorescent reporter plasmids were maintained with 7.5 μg/mL (pCM29) or 10 μg/mL (pEM87) chloramphenicol. For all other experiments, *P. aeruginosa* and *Escherichia coli* were grown in lysogeny broth (LB), and *S. aureus* in tryptic soy broth (TSB; BD Bacto^TM^), unless otherwise specified. Bacteria were grown overnight rotating at 30°C (*E. coli* with pMQ30 background vector) or 37°C for 14 to 16 h. Subcultures were also grown rotating at their respective temperatures.

### Generation of P. aeruginosa deletion mutants

Markerless deletion mutants of genes in PA14 were constructed through homologous recombination as previously described [61]. Briefly, to generate the Δ*lasA* mutants, ∼800 bp upstream and downstream of *lasA* were amplified from WT *P. aeruginosa* PA14 chromosome with primers ASP12_lasA_KO_HindIII_UP_F and ASP13_lasA_KO_UP_R (upstream) and ASP14_lasA_KO_DN_F and ASP15_lasA_KO_SacI_DN_R (downstream) (Table S3) and cloned into pEXG2-Tc with Gibson assembly (New England Biolabs). pEXG2-Tc-Δ*lasA* (Table S2) was introduced into *E. coli* S17 and conjugated with *P. aeruginosa* to perform two-step allelic exchange, and counter selected with 10% sucrose. The Δ*pilA* deletion mutants were generated with the pSMC259-Δ*pilA* (Table S2) following the same protocol. Deletion mutants were confirmed by PCR with primers ASP18_lasA_KO_seq_Fwd and ASP19_lasA_KO_seq_Rev for Δ*lasA*, and oDHL34_pilA-check-F and oDHL35_pilA-check-R for Δ*pilA* (Table S3) for the regions flanking the target genes, followed by Sanger sequencing of the PCR products.

### Integration of mini-Tn7-*mCherry* into *P. aeruginosa* chromosome

The mini-Tn7 element was integrated into *P. aeruginosa att*Tn7 site following a modified protocol from Choi & Schweizer (2006) [62]. Briefly, *P. aeruginosa* overnights were grown in TSB and made electrocompetent with 300 mM sucrose. 300 - 500 ng of pBT277 and the helper plasmid pTNS3 each were simultaneously electroporated into *P. aeruginosa* target strains (not exceeding 5 μL total). To recover, cells were incubated rotating at 37°C for 1.5 h and then plated on TSB agar with 30 μg/mL gentamicin, and left incubating overnight at 37°C.

### Time-lapse microscopy

*P. aeruginosa-*mCherry strains and *S. aureus* pCM29 were grown overnight in M8T and subcultured in fresh M8T (no antibiotic selection) until mid-exponential phase (OD600 of ∼0.3 - 0.8). *P. aeruginosa* and *S. aureus* were standardized to OD600 of 0.010 and 0.020, respectively, in pre-warmed M8T. Agarose pads were prepared by pipetting 550 µL of M8T with 2% molten agarose (Lonza, Cat. no. 50081) into each quadrant of a 4-chamber glass-bottom dish (Cellvis, Cat. no. D35C4-30-1.5-N) and drying uncovered for ∼1.5 h at room temperature, followed by ∼1 h covered with a lid at room temperature, then ∼1.5 h at 37°C. The standardized *P. aeruginosa* and *S. aureus* were mixed 1:1 in a 1.5 mL tube, vortexed, and 0.4 µL were added to one quadrant of a 4-chamber glass-bottom dish before transferring the pads onto the inoculated glass-bottom dish. For monoculture conditions, *S. aureus* or *P. aeruginosa* were mixed with M8T 1:1. Fields of view with *S. aureus* and *P. aeruginosa* single cells positioned approximately 30 to 50 µm apart were selected. Resonant scanning confocal microscopy was employed with a Nikon Eclipse Ti2 A1R and a 60x Plan Apo λ oil objective (1.4 NA). Two Z-stacks (40 - 60 μm) with a 0.12 μm step size, at different XY positions per condition were acquired at 1 h intervals for 18 h, followed by an end time point at ∼24 h. A 488 nm laser (laser power between 3 - 5%, offset −15, and HV – gain between 15 - 25) was used to excite the sGFP produced by *S. aureus* cells, and a 561 nm laser (laser power between 8 - 10%, offset 0, and HV – gain 45) to excite the mCherry in *P. aeruginosa* strains, with a pinhole set between 30 and 39 μm. The phase contrast images were acquired with an Andor Sona camera using a 100x Plan Apo λ oil Ph3 objective (1.45 NA) with 1.5x zoom. All images were saved and analyzed with the Nikon NIS-Elements AR Software.

### *S. aureus* colony edge height and density measurements

*S. aureus* pCM29 and *P. aeruginosa-*mCherry were grown overnight and subcultured in M8T and prepared for imaging in 4-chamber dishes as described above. At approximately 24 h, galvanometer scanning confocal microscopy was employed to image 30 μm Z-stacks (0.12 μm step size) of *S. aureus* colony edges with a 60x Plan Apo λ oil objective (1.4 NA) with 1.5x zoom. The 488 nm laser (laser power 4%, offset −15, and HV – gain 20) with a pinhole of 35 μm was used to excite *S. aureus* GFP cells.

A vertical cross-section crop 15 µm into the colony was performed, and the height on the Z-plane was assessed using the measuring tool for volume projections in Nikon Elements.

The density of *S. aureus* was determined with the BiofilmQ framework [41], by quantifying the biofilm surface roughness and the architecture local density (cell packing) parameters. Segmented microbial volumes were divided into a 3D grid with each node 0.72 μm on a side. Neighborhood cell packing measurements calculated the local biovolume fraction within 6 μm of each segmented bacterial volume within each grid cube. Biofilm surface roughness was calculated by measuring the surface area within 4μm around each grid cube. Each pixel within the heatmaps (Fig. 2D and E) shows the average roughness or cell packing value at every height in the Z-stack at that specific XY coordinate.

### *S. aureus* lactic acid fermentation (P_*ldh1-sgfp*_) quantification

*P. aeruginosa-*mCherry and *S. aureus* pEM87 were grown overnight in M8T. Overnight cultures were subcultured in fresh M8T with 10 μg/mL chloramphenicol used for *S. aureus*. Bacteria were then prepared for imaging under agarose pads as described above. Time-lapse resonant scanning confocal microscopy was performed as described above with a 60x Plan Apo λ oil objective (1.4 NA), for a total of 18 h with 1 h intervals. A 488 nm laser (laser power 6%, offset −20, and HV – gain 30) was used to excite the sGFP in *S. aureus* cells, and a 561 nm laser (laser power 6%, offset 0, and HV – gain 45) to excite the mCherry in *P. aeruginosa* strains, with a pinhole of 39.6 μm.

The Mean Fluorescence Intensity (MFI) from *S. aureus* colonies was measured in the Nikon Elements software using the 3D Measurements tool. This measurement takes into consideration the volume of the colony and is calculated with the following equation: 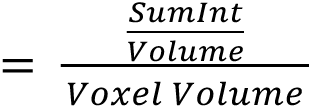. The threshold used for *S. aureus* colonies was the following: 150 - 4095 for brighter colonies (*S. aureus* in coculture with WT or Δ*pilA P. aeruginosa*) or 50 - 4095 for dimmer colonies (*S. aureus* in coculture with Δ*pqsL* or Δ*pqsL* Δ*pilA P. aeruginosa*).

### Arabinose-inducible genetic complementation of *pqsL* and *pilA*

S. aureus pEM87 and P. aeruginosa strains were grown overnight in M8T. P. aeruginosa strains harboring the pMQ72-P_*araBAD*_ empty vector or pMQ72-P_*araBAD-pqsL*_ (Table S2) were cultured with 30 μg/mL gentamicin in the overnights and subcultures. No antibiotic selection was used for the ΔpilA complementation strains which contain the chromosomal attTn7 arabinose-inducible system (Table S1). Strains were subcultured in M8T until mid-exponential phase with 0.4% arabinose in the P. aeruginosa cultures. Cells were then prepared, imaged, and analyzed as described above using the same settings as the previous section (S. aureus P_*ldh1-sgfp*_), with the addition of 0.4% arabinose to the agarose pads to induce P_*araBAD-pqsL*_ or -_*pilA*_.

### Artificial sputum media assay

*P. aeruginosa-*mCherry and *S. aureus* pCM29 were grown overnight in M8T. The overnight liquid cultures were subcultured in fresh M8T (no antibiotic selection) and grown rotating until mid-exponential phase. Cultures were normalized to an OD600 of 0.2 in M8T and then washed 1X in PBS. The artificial sputum media (ASM) was prepared as previously described [5]. The washed cultures were then inoculated into ASM at an OD600 of 0.025 in mono- or coculture in a total volume of 500 μL in a 1.5 mL microcentrifuge tube. The tubes were mixed by vortexing, and then 200 μL were inoculated into each quadrant of a 4-chamber glass-bottom 35 mm dish. The bacteria were grown statically for ∼24 h inside the microscope’s incubation chamber at 37°C with 90% relative humidity controlled by the Okolab Humidity Controller. At ∼22 h, two 50 μm Z-stacks at different XY positions per condition were acquired with an inverted Nikon Eclipse Ti2 A1R Resonant Scanning Confocal Microscope, using a CFI SR HP Plan Apochromat Lambda S 100XC Sil objective (1.35 NA) with a 56.19 μm pinhole and 1024 x 1024 pixels. A 488 nm laser (laser power 5, offset −10, and HV – gain 10) was used to excite the sGFP in *S. aureus* cells, and a 561 nm laser (laser power 7, offset −10, and HV – gain 40) to excite the mCherry in *P. aeruginosa* strains.

For the experiments where *P. aeruginosa* was added to *S. aureus* pre-formed biofilms, *S. aureus* was prepared the same way as described above and incubated statically in monoculture for 4 - 5 h. *P. aeruginosa* strains at a final concentration of OD600 0.03 were then added to *S. aureus* and incubated for an additional 24 h. Then, 50 μm image stacks were acquired at ∼29 h.

The images were saved and analyzed with the Nikon Elements software. The representative images in Figs. 4 and 5 were deconvolved and are projected as a volume view with the Maximum Intensity Projection blending.

*S. aureus* and *P. aeruginosa* growth was assessed at ∼24 h (when *P. aeruginosa* was coinoculated with *S. aureus* at T0) or ∼29 h (when *P. aeruginosa* was added to pre-formed *S. aureus* biofilms) by plating the serial dilutions on selective media (Mannitol Salt Agar (MSA) and *Pseudomonas* Isolation Agar (PIA), respectively). To harvest the bacteria from the microscopy dishes, 0.2% Triton X-100 was added to the cultures and incubated at room temperature for 10 - 20 m. Then, the bacteria were scrapped with a pipette tip and all the volume per quadrant was transferred to a 1.5 mL centrifuge tube. The tubes were thoroughly mixed by vortexing prior to performing the serial dilutions.

### *S. aureus* cell disruption measurement

The end-time point Z-stack images from the artificial sputum data (Fig. 5 and Supplementary Fig. 4) were analyzed in the Nikon Elements analysis software. Using the 3D Measurements tool, the volume of the bottom four Z planes (i.e., *S. aureus* cells attached to the coverslip) was divided by the volume of the rest (top part) of the *S. aureus* biofilm.

### *P. aeruginosa* supernatant collection

*P. aeruginosa* strains were grown overnight in LB. The cultures were standardized to OD600 3.0 in LB and spun for 5 m at 15 krpm. The supernatants were filter sterilized with a PES 0.22 μm filter.

### *S. aureus* lysis assay with *P. aeruginosa* supernatant

*S. aureus* lysis was assessed by modifying a previously published method [63]. *S. aureus* and *P. aeruginosa* were grown overnight in TSB or LB, respectively. *P. aeruginosa* cell-free supernatants were collected as described above. *S. aureus* was subcultured in TSB starting at OD600 0.1 and grown for 3 h. All the volume from the cultures (5 mL) was centrifuged for 10 m at 4 krpm, washed twice in the same volume of cold (4°C) sterile distilled water, and resuspended in buffer (50 mM Tris-HCl (pH 7.2) with 0.05% Triton X-100). The bacteria were then standardized to OD600 1.0 in buffer and 500 μL of *P. aeruginosa* supernatant (OD600 3.0) were added to triplicate plastic cuvettes followed by 500 μL of *S. aureus* in buffer (OD600 1.0). For the *S. aureus* control, 500 μL of buffer plus 500 μL of *S. aureus* cells were used. Cuvettes were covered with parafilm, mixed by inversion, and the initial OD580 measurement was acquired with a spectrophotometer. Then, the samples were incubated shaking (225 rpm) at 30°C, and measurements were taken every 30 m for a total of 2.5 h.

### *S. aureus* growth curve with *P. aeruginosa* supernatant

*S. aureus* and *P. aeruginosa* were grown overnight in TSB or LB, respectively. *S. aureus* was subcultured in TSB until mid-exponential phase. *P. aeruginosa* supernatants were collected as described above. *S. aureus* cells were normalized to OD600 0.02 in TSB and mixed 1:1 with *P. aeruginosa* supernatant (collected from cultures at OD600 3.0) in a 1.5 mL microcentrifuge tube, for a final concentration of *S. aureus* at OD600 0.01 and *P. aeruginosa* supernatant at OD600 1.5. For monoculture *S. aureus*, cells were mixed 1:1 with LB, since the *P. aeruginosa* supernatants were derived from cultures grown in LB. The tubes were thoroughly mixed, and 200 μL were added to triplicate wells of a clear flat bottom 96-well plate. The plate was covered with a gas-permeable membrane and OD600 measurements were taken every 15 m on a Tecan infinite M200 plate reader for 18 h total. The plate was incubated statically at 37°C and was shaken for 30 s before each measurement.

## ACKNOWLEDGEMENTS

We thank Dr. George O’Toole’s lab and Dr. Timothy Yahr for generously providing bacterial strains and plasmids. We also thank Dr. Michael J. Gebhardt and members of the Limoli lab for their insightful feedback on the manuscript.

This work was supported by funding from the Cystic Fibrosis Foundation: CFF Postdoc-to-Faculty Transition Award LIMOLI18F5 (DHL), CFF RDP Junior Faculty Recruitment Award LIMOLI19R3 (DHL), and funding from the National Institutes of Health: Maximizing Investigators’ Research Award R35GM142760 (DHL), 1R35GM151158-01 (CDN), the Molecular & Cellular Biology of the Lung Training Grant 5T32HL007638-34 (ASP), and the T32 Training Grant AI007519 (JBW). Additional support was provided by the John H. Copenhaver Jr. Fellowship (JBW), the GAANN Fellowship (JBW), the Human Frontier Science Program RGY0077/2020 (CDN), the Simons Foundation 826672 (CDN), and by the National Science Foundation IOS grant 2017879 (CDN).

## COMPETING INTERESTS

The authors have declared no competing interests.

## SUPPLEMENTARY MATERIAL

**Supplementary Figure 1.**
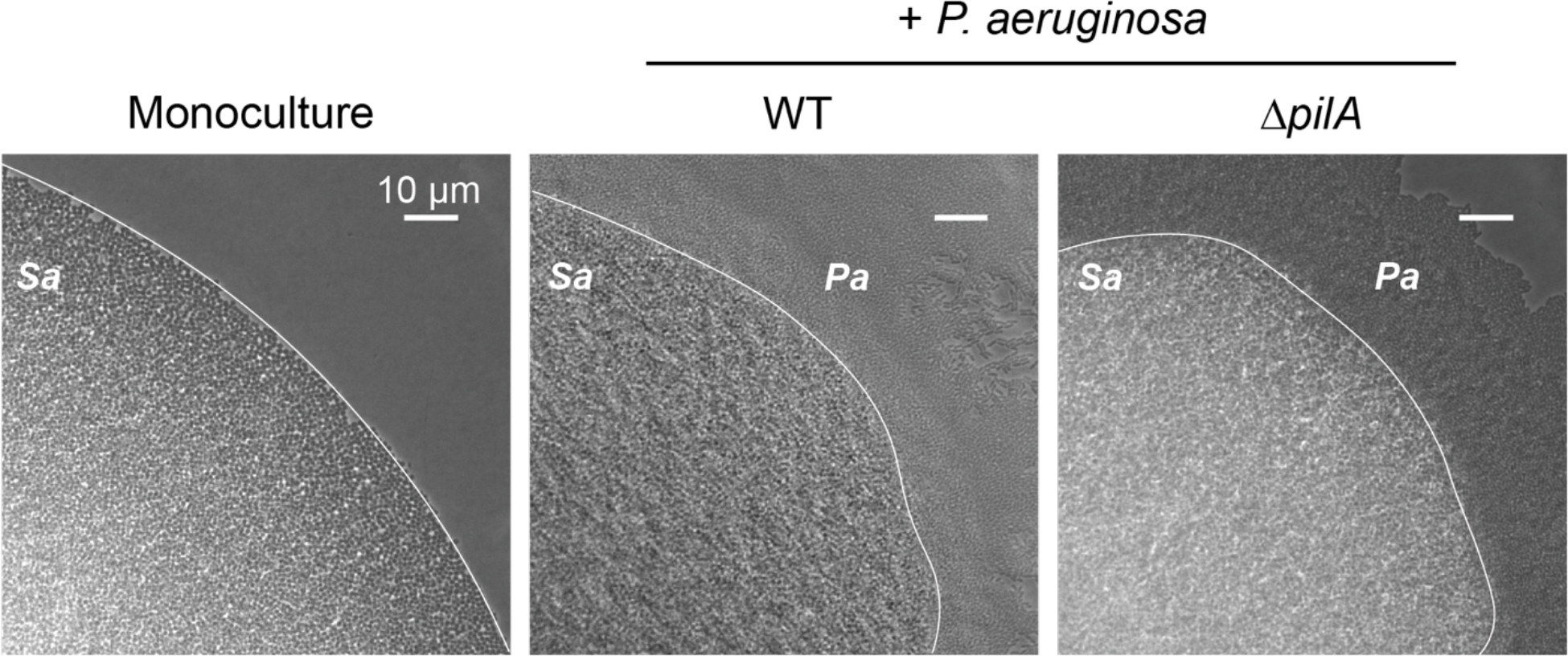
*P. aeruginosa* cells surround *S. aureus* colonies at the end time point. Representative phase contrast micrographs of *S. aureus* in mono- or coculture with WT or Δ*pilA P. aeruginosa* at ∼24 h. The white line traces *S. aureus* colony border.

**Supplementary Figure 2.**
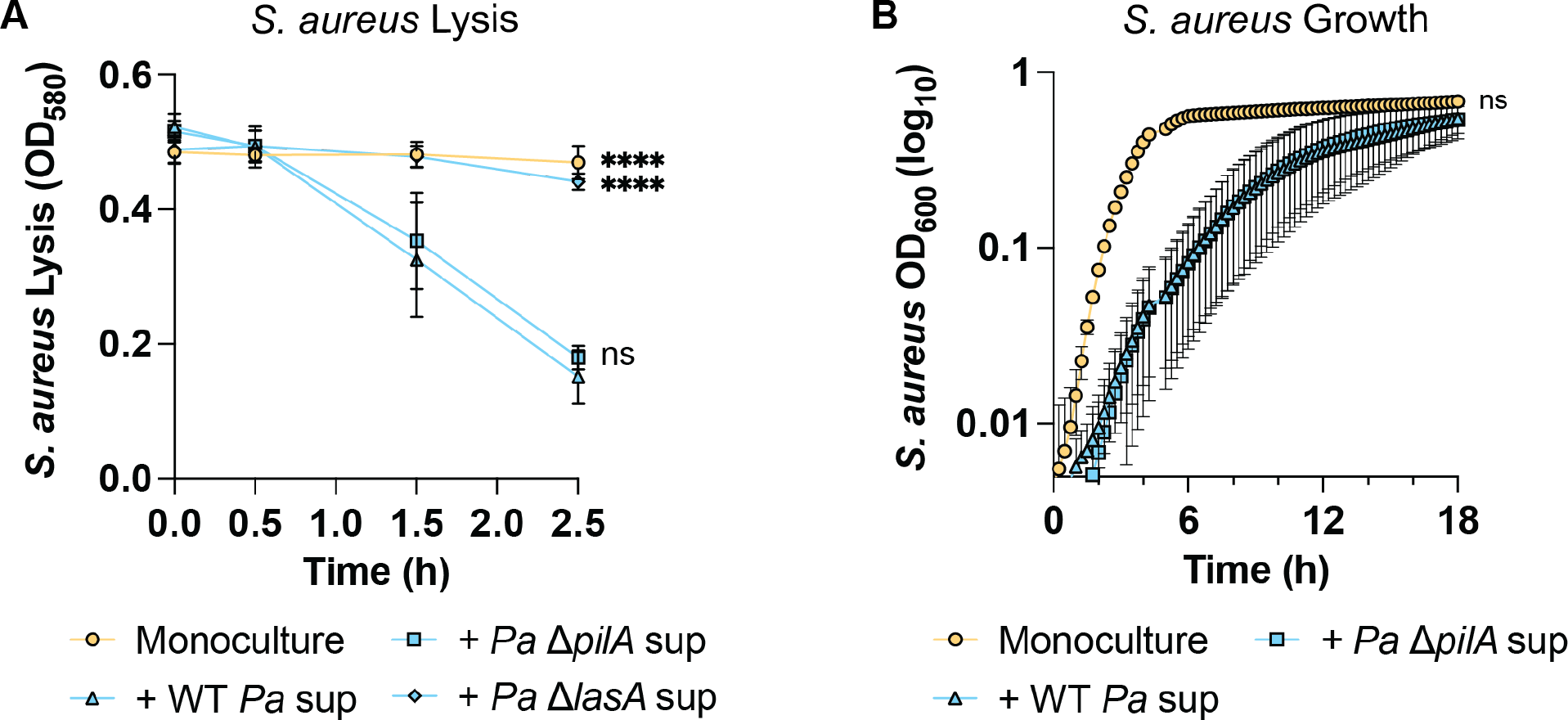
Exoproducts from Δ*pilA* lyse and inhibit *S. aureus* to the same levels as WT *P. aeruginosa* factors. *S. aureus* lysis (**A**) or growth (**B**) in the presence of WT, Δ*pilA*, or Δ*lasA P. aeruginosa* supernatant. Three (**A**) or four (**B**) biological replicates with three technical replicates each were performed. Statistical significance was determined by one-way ANOVA followed by Dunnett’s Multiple Comparisons Test at 2.5 h (**A**), or a two-way ANOVA followed by Tukey’s Multiple Comparisons Test at every time point (**B**). No difference was found between the + WT *Pa* sup and the + *Pa* Δ*pilA* sup conditions in **B** at any time point. n.s. = not significant; **** = *P* < 0.0001.

**Supplementary Figure 3.**
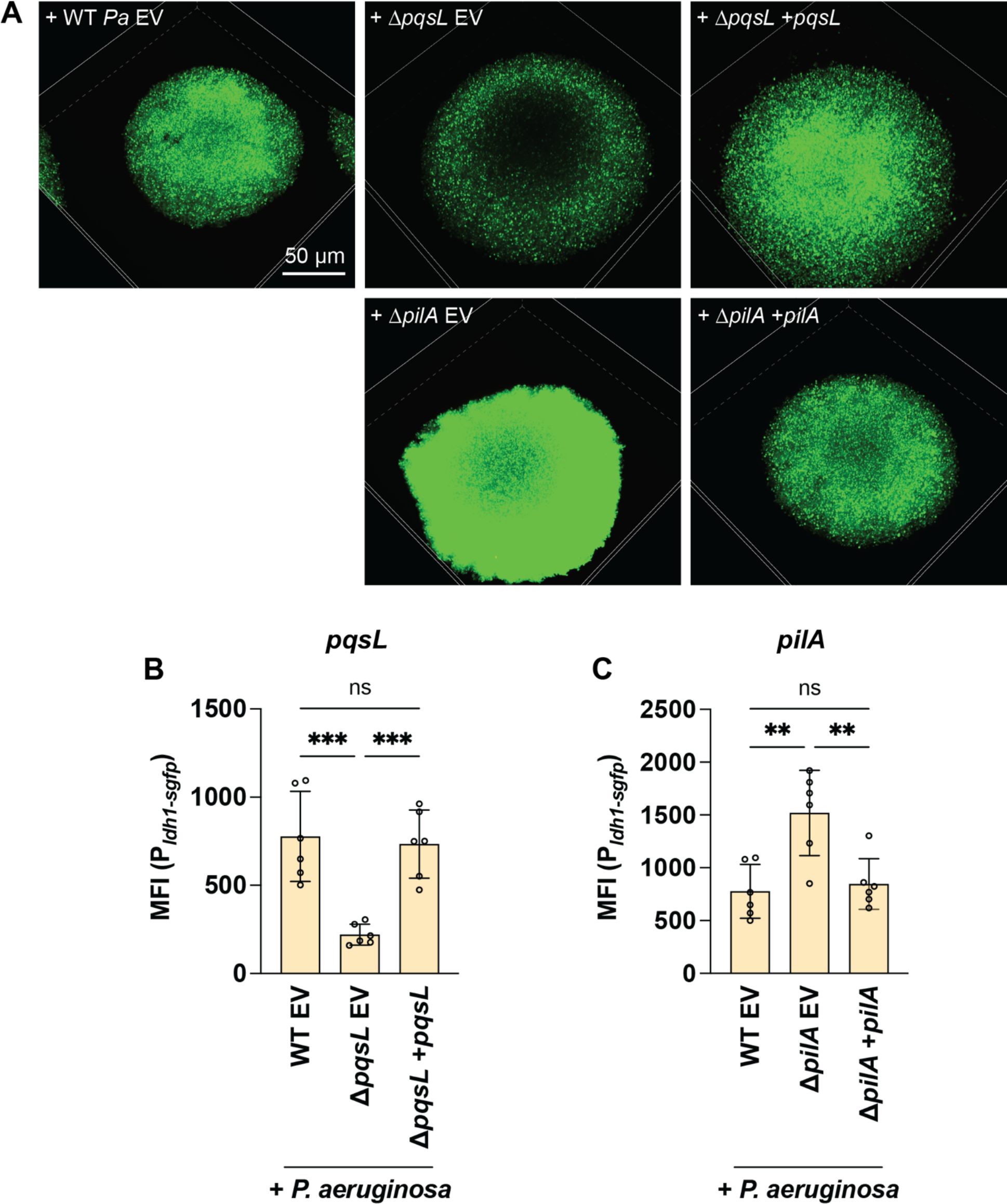
Genetic complementation of *pqsL* or *pilA*. *S. aureus* lactate fermentation (P_*ldh1-sgfp*_) was measured in the presence of the indicated *P. aeruginosa* strains. **A.** Representative resonant scanning confocal micrographs of *S. aureus* fermentation in coculture with WT *P. aeruginosa* pMQ72 P_*araBAD*_ empty vector (EV), Δ*pqsL* pMQ72 EV, Δ*pqsL* pMQ72-P_*araBAD-pqsL*_, Δ*pilA attTn7::*EV, or Δ*pilA attTn7::*P_*araBAD-pilA*_ at t = 18 h. **B & C.** The mean fluorescence intensity (MFI) of *S. aureus* colonies was quantified in the presence of *P. aeruginosa pqsL* (**B**) or *pilA* (**C**) complementation strains at 18 h for three biological replicates with two technical replicates each, and the mean and standard deviation are shown. Each data point represents one technical replicate. Statistical significance was determined by one-way ANOVA followed by Dunnett’s Multiple Comparisons Test. n.s. = not significant; ** = *P* < 0.01; *** = *P* < 0.001.

**Supplementary Figure 4.**
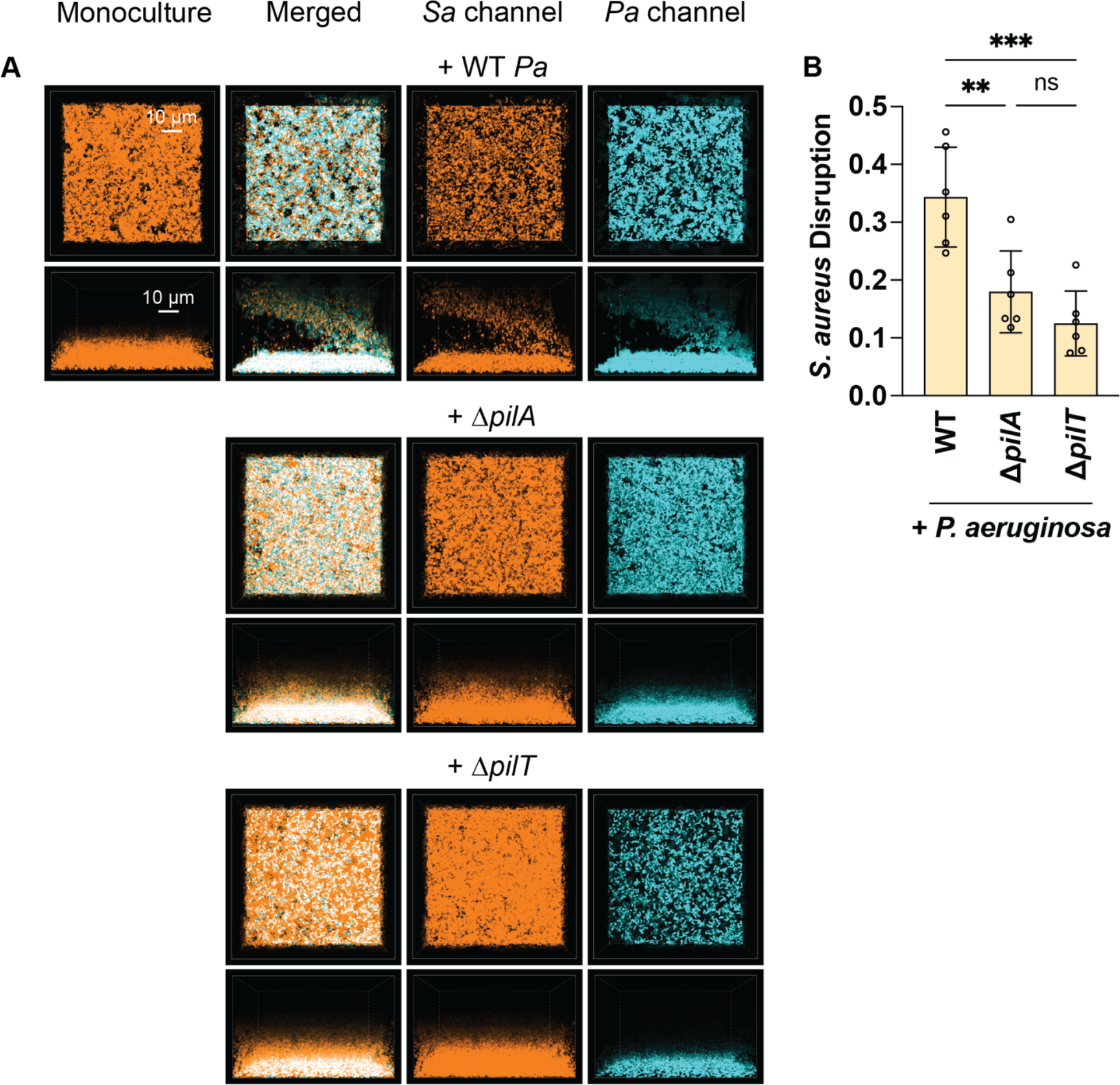
*P. aeruginosa* type IV pili motility is necessary for disrupting pre-formed *S. aureus* biofilms. A. Representative resonant scanning confocal micrographs of *S. aureus* and *P. aeruginosa* in artificial sputum media. WT *S. aureus* (pseudocolored orange) in monoculture or in coculture with *P. aeruginosa* (pseudocolored cyan; WT, Δ*pilA*, or Δ*pilT*) t ∼29 h. **B.** *S. aureus* biofilm disruption quantification (*Sa* volume at the coverslip / *Sa* volume at the top) at t ∼29 h. Three biological replicates with two technical replicates each were analyzed. The mean and standard deviation are shown. Each data point represents one technical replicate. Statistical significance was determined by one-way ANOVA followed by Dunnett’s Multiple Comparisons Test. n.s. = not significant; ** = *P* < 0.01; *** = *P* < 0.001.

**Supplementary Table 1.** Bacterial strains used in this study.

| Strain | Alternative Strain Names | Genotype or Description | Reference or Source |
| --- | --- | --- | --- |
| *Escherichia coli* |  |  |  |
| NEB® 5-alpha competent *E. coli* |  |  | New England Biolabs |
| S17 λpir | ECDHL23 | *pro*, *res*<sup>-</sup> *hsdR17* (*rK*<sup>-</sup> *mK*<sup>+</sup>) *recA*<sup>-</sup> with an integrated *RP4-2-Tc::Mu-Km::Tn7*, T<sub>p</sub><sup>r</sup> | [1] |
| *Pseudomonas aeruginosa* |  |  |  |
| PA14 | SMC232 | Nonmucoid | [2] |
| PA14 Δ*pilA* | SMC3782 | | [3] |
| PA14 Δ*lasA* | PADHL447 | | This study |
| PA14 Δ*pqsL* Δ*pilA* | PADHL572 | | This study |
| PA14 mCherry | PADHL441 | *attTn7::miniTn72.1-Gm-GW:P<sub>A1/04/03-mCherry</sub>*; Gm<sup>r</sup>. | This study |
| PA14 Δ*pilA* mCherry | PADHL439 | *attTn7::miniTn72.1-Gm-GW:P<sub>A1/04/03-mCherry</sub>*; Gm<sup>r</sup>. | This study |
| PA14 Δ*pilT* mCherry | PADHL542 | *attTn7::miniTn72.1-Gm-GW:P<sub>A1/04/03-mCherry</sub>*; Gm<sup>r</sup>. | This study |
| PA14 Δ*lasA* mCherry | PADHL461 | *attTn7::miniTn72.1-Gm-GW:P<sub>A1/04/03-mCherry</sub>*; Gm<sup>r</sup>. | This study |
| PA14 Δ*pqsL* mCherry | PADHL442 | *attTn7::miniTn72.1-Gm-GW:P<sub>A1/04/03-mCherry</sub>*; Gm<sup>r</sup>. | This study |
| PA14 Δ*pqsL* Δ*pvdA* Δ*pchE* mCherry | PADHL440 | *attTn7::miniTn72.1-Gm-GW:P<sub>A1/04/03-mCherry</sub>*; Gm<sup>r</sup>. | This study |
| PA14 Δ*pqsL* Δ*pvdA* Δ*pchE* Δ*lasA* mCherry | PADHL463 | *attTn7::miniTn72.1-Gm-GW:P<sub>A1/04/03-mCherry</sub>*; Gm<sup>r</sup>. | This study |
| PA14 Δ*pqsL* Δ*pvdA* Δ*pchE* Δ*lasA* Δ*pilA* mCherry | PADHL544 | *attTn7::miniTn72.1-Gm-GW:P<sub>A1/04/03-mCherry</sub>*; Gm<sup>r</sup>. | This study |
| PA14 pMQ72-*P<sub>araBAD</sub>* empty vector | PADHL865 | Gm<sup>r</sup> | This study |
| PA14 $\Delta pqsL$ pMQ72-$P_{araBAD}$ empty vector | SMC6232 | Gm<sup>r</sup> | [4] |
| PA14 $\Delta pqsL$ pMQ72-$P_{araBAD-pqsL}$ | SMC6231 | Gm<sup>r</sup> | [4] |
| PA14 $\Delta pilA$ attTn7::$P_{araBAD}$ empty vector | SMC7456 | Gm<sup>r</sup> | [5] |
| PA14 $\Delta pilA$ attTn7::$P_{araBAD-pilA}$ | SMC7457 | Gm<sup>r</sup> | [5] |
| ***S. aureus*** |  |  |  |
| USA300 JE2 | SADHL05 | USA300 CA-Methicillin resistant strain<br>LAC without plasmids | [6] |
| JE2 pCM29 | SADHL07 | Cm<sup>r</sup> | [7] |
| JE2 pEM87 | SADHL167 | Cm<sup>r</sup> | [8] |

Cm, chloramphenicol; Gm, gentamicin

**Supplementary Table 2.**
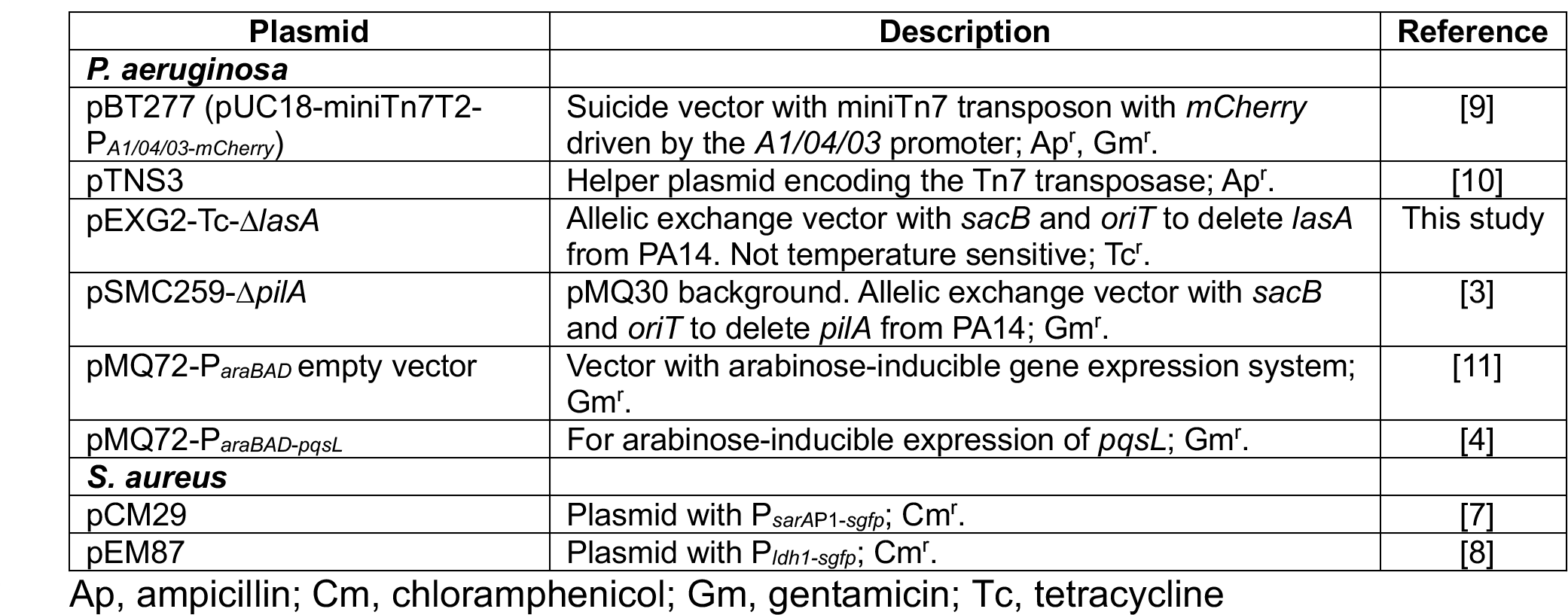
Plasmids used in this study.

| Plasmid | Description | Reference |
| --- | --- | --- |
| ***P. aeruginosa*** |  |  |
| pBT277 (pUC18-miniTn7T2-$P_{A1/04/03-mCherry}$) | Suicide vector with miniTn7 transposon with *mCherry* driven by the *A1/04/03* promoter; Ap<sup>r</sup>, Gm<sup>r</sup>. | [9] |
| pTNS3 | Helper plasmid encoding the Tn7 transposase; Ap<sup>r</sup>. | [10] |
| pEXG2-Tc-$\Delta lasA$ | Allelic exchange vector with *sacB* and *oriT* to delete *lasA* from PA14. Not temperature sensitive; Tc<sup>r</sup>. | This study |
| pSMC259-$\Delta pilA$ | pMQ30 background. Allelic exchange vector with *sacB* and *oriT* to delete *pilA* from PA14; Gm<sup>r</sup>. | [3] |
| pMQ72-$P_{araBAD}$ empty vector | Vector with arabinose-inducible gene expression system; Gm<sup>r</sup>. | [11] |
| pMQ72-$P_{araBAD-pqsL}$ | For arabinose-inducible expression of *pqsL*; Gm<sup>r</sup>. | [4] |
| ***S. aureus*** |  |  |
| pCM29 | Plasmid with $P_{sarAP1-sgfp}$; Cm<sup>r</sup>. | [7] |
| pEM87 | Plasmid with $P_{ldh1-sgfp}$; Cm<sup>r</sup>. | [8] |

Ap, ampicillin; Cm, chloramphenicol; Gm, gentamicin; Tc, tetracycline

**Supplementary Table 3.**
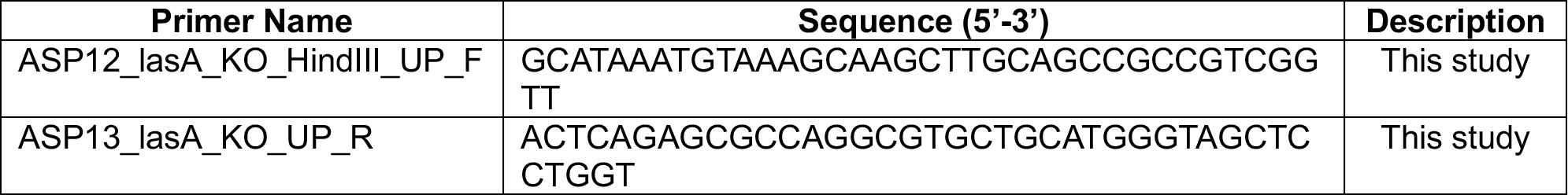
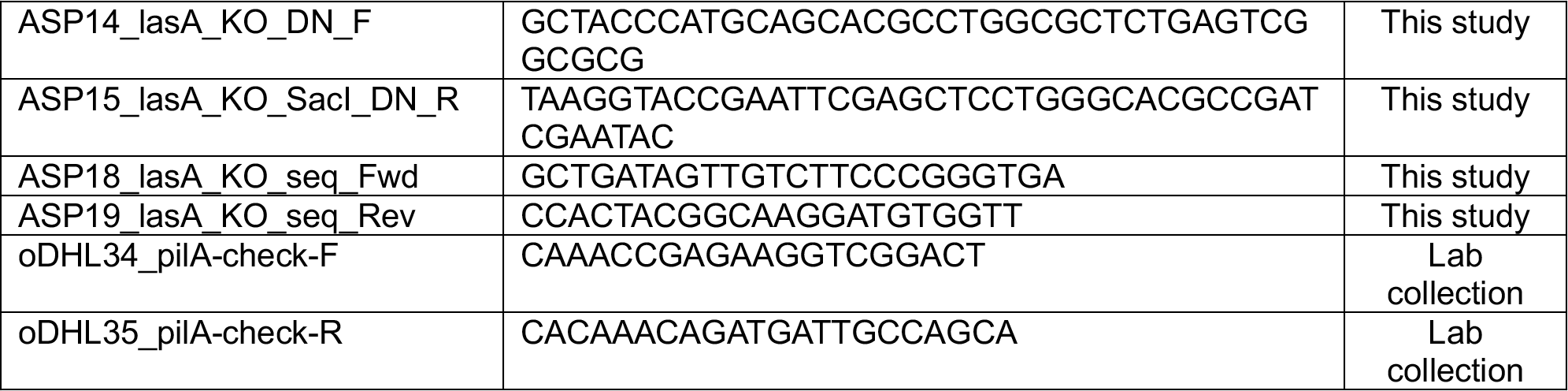
Primers used in this study.

